# Disturbance Triggers Non-Linear Microbe-Environment Feedbacks

**DOI:** 10.1101/2020.09.30.314328

**Authors:** Aditi Sengupta, Sarah J. Fansler, Rosalie K. Chu, Robert E. Danczak, Vanessa A. Garayburu-Caruso, Lupita Renteria, Hyun-Seob Song, Jason Toyoda, Jacqueline Wells, James C. Stegen

## Abstract

Conceptual frameworks linking microbial community membership, properties, and processes with the environment and emergent function have been proposed but remain untested. Here we refine and test a recent conceptual framework using hyporheic zone sediments exposed to wetting/drying transitions. Throughout the system we found threshold-like responses to the duration of desiccation. Membership of the putatively active community--but not the whole community--responded due to enhanced deterministic selection (an emergent community property). Concurrently, the thermodynamic properties of organic matter became less favorable for oxidation (an environmental component) and respiration decreased (a microbial process). While these responses were step functions of desiccation, we observed continuous monotonic relationships among community assembly, respiration, and organic matter thermodynamics. Placing the results in context of our conceptual framework points to previously unrecognized internal feedbacks that are initiated by disturbance, mediated by thermodynamics, and that cause the impacts of disturbance to be dependent on the history of disturbance.

## Introduction

Given the influence of microbes over ecosystem function, deeper knowledge of microbe-environment relationships is needed to improve ecosystem models (*1*). In turn, there is strong interest in quantifying and predicting microbe-environment relationships such as defining microbial life history strategies as traits in ecosystem models (*2*), assessing microbial biomass stoichiometry distributions in response to changing resource environments (*3*), and evaluating the extent of microbial adaptation to changing environments and their role in biogeochemical processes (*4*). To enhance and synthesize understanding of microbe-environment interactions, it is useful to develop conceptual frameworks based on linkages among microbial characteristics and ecosystem processes. Previous work has used such frameworks to improve mechanistic representation and predictive capacity of microbe-environment interactions in ecosystem models (*5*).

A recently developed framework by Hall et al. (*6*) poses a series of concepts that collectively define the intersection between microbial and ecosystem ecology. Their framework draws attention to causal relationships between microbial characteristics, environmental dynamics, and cumulative ecosystems processes with the potential to incorporate relevant mechanistic links into predictive ecosystem models. A powerful element of the Hall et al. (*6*) framework is that it applies to diverse systems spanning natural (*7*), host-associated (*8*), and built (*9*) environments as well as across spatiotemporal scales (*10*). While potentially very useful, the Hall et al. (*6*) framework has seen little direct use in terms of explicitly defining and evaluating the linkages within specific study systems (but see *3*). To make full use of and continually improve the framework, it is necessary to consider different realizations and interpretations of the proposed linkages. In the following paragraphs we detail a modified interpretation of the framework (**Fig. 1**) to enable its application to microbial communities and biogeochemistry associated with hyporheic zone sediments experiencing hydrological disturbance. In turn, we use data from a controlled laboratory experiment to evaluate key linkages within the modified framework.

**Figure 1.**
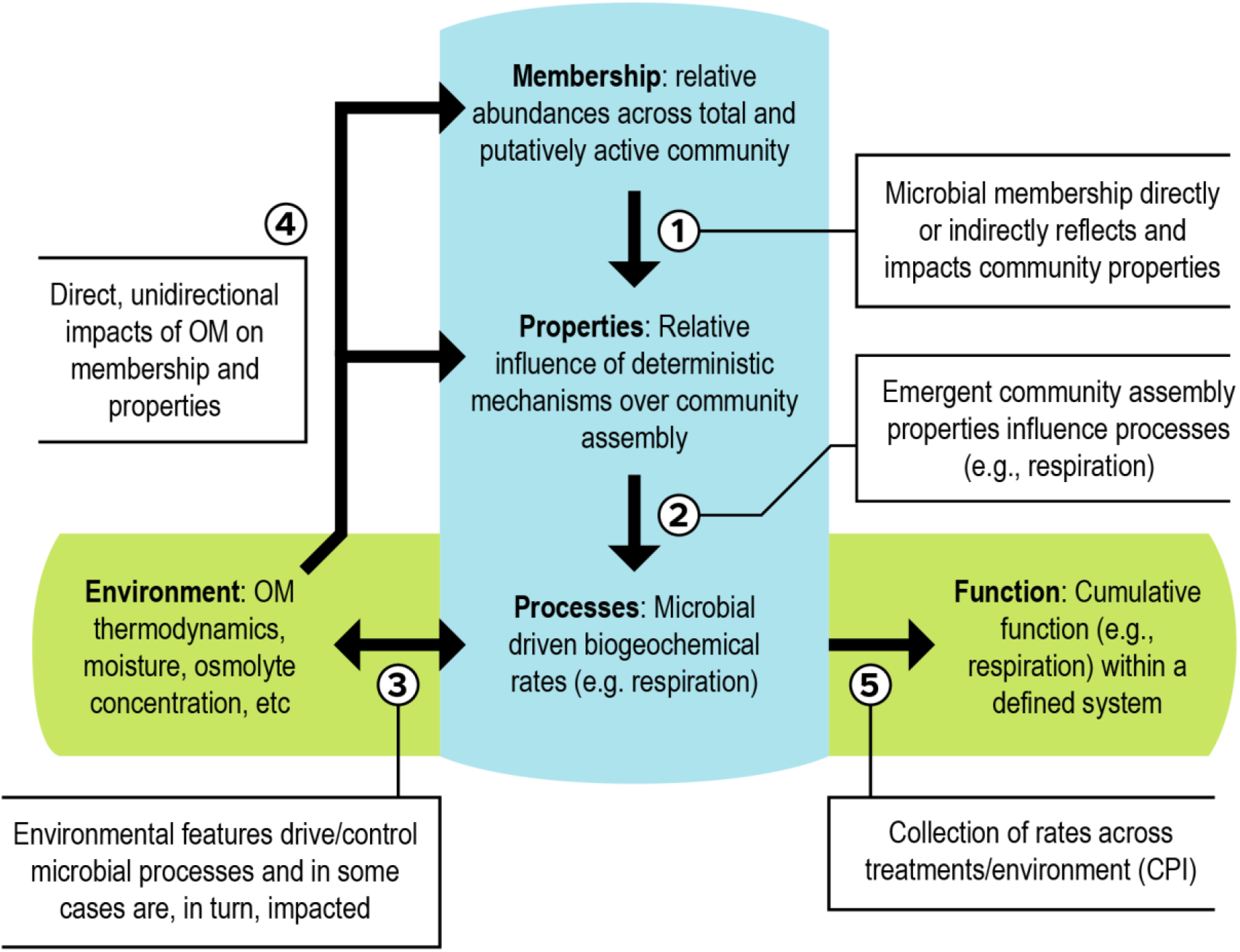
Integrated conceptual framework. The conceptual figure (modified from Hall et al. (*6*)) details relationships (indicated by numbered arrows) between cumulative properties of the microbial community (*e.g*., microbial membership, community assembly properties, biogeochemical rates), environmental features (*e.g.*, organic matter thermodynamics), and emergent ecosystem function

As in Hall et al. (*6*) we consider microbial membership to be directly influenced by environmental conditions (arrow 4, Fig. 1) and to underlie community-level properties (arrow 1, Fig. 1). Determining microbial membership is relatively straightforward, and uses culture-independent (*11–14*) and culture-dependent (*15*) techniques. Sequence-based assays using phylogenetic markers are routine, with DNA-based (total community members) and RNA-based (putatively active community members) (*16–21*) approaches providing the foundation to study community properties.

While community membership is relatively straightforward, the identification of community properties that are relevant to a given system and function is open to broader interpretation. Here we propose using the relative influences of deterministic and stochastic community assembly (*22*) as emergent properties of microbial communities that have implications for biogeochemical function (*23*) including in ecosystems experiencing environmental disturbance. Deterministic mechanisms are associated with systematic differences in reproductive success imposed by the biotic and/or abiotic environment, while stochastic mechanisms are associated with passive spatial movements of organisms and birth/death events that are not due to systematic differences across taxa in reproductive success (*24, 25*). The relative contributions of determinism and stochasticity can be inferred by coupling microbial community membership and phylogeny to ecological null models (*22, 25–27*).

We propose the relative contributions of determinism and stochasticity to be an emergent property that is greater than the sum of the individual components (i.e., taxa), and that is complementary to the community properties proposed by Hall et al. (*6*), such as biomass and gene expression. Note that here we conceptualize stochastic and deterministic events as ecological community assembly processes and that the relative influences of these ecological processes as an emergent community property. The ecological processes of community assembly are distinct from ‘microbial processes’ associated with biogeochemical reactions. As an emergent property, the relative influence of determinism and stochasticity is the result of complex biotic and abiotic interactions (*28*) and also shapes cumulative microbial processes that impact ecosystem biogeochemical functions (*23*) (arrow 2, Fig. 1). For example, a stronger influence of determinism over community assembly is hypothesized to cause higher respiration rates (a microbial processes) due to a larger contribution of well-adapted taxa (*23*).

Analyses of microbial community assembly have been widely employed across environments including soil (*24, 29–32*), sediment (*26, 33–35*), marine (*36, 37*), riverine (*38*), gut (*39*), and engineered (*40, 41*) systems. Previous work has focused primarily on using DNA-derived membership and phylogenetic data to study whole-community assembly. In contrast, recent studies have also used an RNA-based approach to study the relative influence of stochasticity over the assembly of the putatively active portion of microbial communities (*31, 42*). This RNA-based approach is complementary to the DNA-based approach and may provide additional insights into shorter-term dynamic linkages between emergent community properties and microbial processes. Such linkages have not, however, been previously evaluated.

The Hall et al. (*6*) framework proposes that microbial processes (e.g., respiration rate) are influenced by both microbial community properties and environmental factors. Here we propose a revision of this structure that includes bidirectional links between the environment and microbial processes (arrow 3, Fig. 1). Such bi-directional links between microbes and their environment are common (*43–46*), and in hyporheic zone sediments may be particularly tied to thermodynamic properties of organic matter (*47*) and influenced by hydrological disturbances that are common in such environments. For example, preferential use OM by microbial communities has potential to alter the thermodynamic properties of organic matter pools (*33*). This microbe-driven shift in the environment could then feedback to impact microbial metabolism due to the strong influence of OM thermodynamics on biogeochemical rates (*34, 48–50*). Bi-directional feedback between environmental factors and microbial processes is, therefore, likely important to the link between microbial communities and ecosystem function. Fundamental knowledge of these feedbacks and how they are modulated by hydrologic disturbances in hyporheic zone sediments is largely unknown, however.

Within ecosystems, mechanistic associations between environmental factors, microbial properties, and microbial processes underlie spatial and temporal distributions of biogeochemical rates (Fig. 1 arrow 5). The resulting distributions (e.g., of respiration rates) define cumulative system function and can be used to understand key phenomena such as biogeochemical hot spots and hot moments (*51*). Developing concepts and models to predict the influences of biogeochemical hot spots/moments is a major outstanding challenge. To facilitate progress, Bernhardt et al. (*52*) proposed grouping hot spots/moments into the concept of ecosystem control points that exert a disproportionate influence on ecosystem function.

While not called out explicitly in Bernhardt et al. (*52*), the control point concept is based on the distribution of biogeochemical rates through space and/or time. Focusing on the shape of rate distributions allows the notion of control points to be extended to the concept of control point influence (CPI; Fig. 1 arrow 5). The CPI is a quantitative measurement of the contribution of elevated biogeochemical rates in space and/or time to the net aggregated rate within a defined system (*53*). While proposed conceptually and studied via simulation in Arora et al. (*53*), empirical measurements of CPI are lacking. More generally, incorporating CPI into a modified version of the Hall et al. (*6*) framework (Fig. 1), provides an integrated conceptualization for how environmental factors, microbial properties, and microbial processes contribute to emergent system function.

Some elements of our modified framework (Fig. 1) are generalizable across systems (e.g., CPI), while others (e.g., OM thermodynamics) may have different levels of relevance across different ecosystem types. Here we aim to generate fundamental knowledge of the linkages between microbial community and ecosystem function, as well as reveal how hydrologic disturbance may modify these linkages. We do so by studying the modified conceptual framework in context of variably inundated hyporheic zone sediments exposed to different drying/wetting dynamics. Hyporheic zones are biogeochemical active subsurface domains in river corridors through which surface water flows and can mix with groundwater (*51, 52, 54*). These zones can have disproportionate biogeochemical impacts on river corridors (*54–58*). Within variably inundated streams (*59, 60*), hyporheic zones experience extreme changes in environmental conditions, but the consequences of this variability for microbe-ecosystem linkages is poorly known.

To mimic natural disturbances we subjected sediments to wetting-drying transitions and focused on a series of analyses tied to our modified framework. We specifically evaluated relationships between: (*i*) the relative influence of stochastic assembly as a community property and respiration rates as a microbial process (Fig. 1 arrow 2), and (*ii*) environmental features and both microbial properties (Fig. 1 arrow 4) and processes (Fig. 1 arrow 3) that underlie aggregate system function (Fig. 1 arrow 5). We hypothesized that: (*i*) Stronger influences of determinism result in well-adapted microbes that will generate higher respiration rates, (*ii*) Longer duration in an inundated state will result in greater influences of stochastic assembly--due to weaker ecological selection--and lower respiration rates following re-inundation due to relatively consistent abiotic conditions (*61*), and (*iii*) Microbial processes are facilitated by OM that is thermodynamically more favorable for oxidation, leading to an association between respiration rates and OM thermodynamics.

## Methods and Materials

### Study site and sediment collection

Hyporheic sediments were collected from the Columbia River shoreline (approximately 46.372411°N, 119.271695°W) in eastern Washington State (Fig S1) (*34, 62–66*) within the Hanford Site on January 14, 2019 at 9 am Pacific Standard Time. Samples were aseptically collected to a depth of 10 cm at five sub-sampling locations within a meter to form a composite sample that was sieved on site through a 2 mm sieve into a clean glass beaker. Sieved sediment was stored on blue ice for 30 min while being transported back to the laboratory. Once back at the laboratory sediment was stored at 4 °C until processing into incubation vials (see below).

The sediments were subjected to increasing temporal environmental variance (as a function of periodic wetting and drying transitions) and evaluated for associations between microbial membership, microbial properties, microbial community assembly, OM chemistry, absolute respiration rate (represented in this study as O_2_ consumption rates), and cumulative respiration rates represented as CPI. Aerobic respiration was chosen as the biogeochemical process since it influences global-scale energy and material fluxes (*67*), and because the hyporheic zone within the field system is predominantly aerobic (*63, 68*). Detailed experimental design and methods are provided in the following paragraphs.

### Experimental design

Sediments used in the batch reactors were sourced from one homogenized sediment pool. In turn, 10 g of sediment from the homogenized pool was added to each reactor vial. The sample set was then split into two groups, one inundated and the other allowed to desiccate. These conditions were maintained for 23 days prior to the start of dynamic moisture manipulation, from January 29th to February 21st, 2019. This initial ‘preconditioning’ period was used to avoid measuring the immediate impacts of sampling disturbance and to allow time for desiccation. All replicate reactors were maintained in the dark, shaking at 100 rmp, at 21 °C and were covered with a gas permeable Breathe-Easy (Milli-Pore Sigma, Burlington, MA) membrane that allowed for gas exchange and drying. After the preconditioning period in which sediments were consistently inundated or allowed to continuously desiccate, the transition regimes (**Fig. 2**) were applied to the reactors starting on February 22nd, 2019. We refer to the time from Feb 22nd, 2019 onwards as the ‘transitions period’ for all treatments, even though some did not experience transitions between being inundated and dry. Each treatment had 6-7 replicates (detailed below). These regimes were designed around the number of wet/dry transitions experienced by sediments within a given treatment. Treatment regimes also caused variation in the cumulative number of days sediments were in a drying state. We imposed six different experimental treatment regimes (**Fig. 2**) as follows:

- 0 Transitions and 0 days of desiccation: Sediments were maintained at field moisture conditions for the preconditioning and transitions periods. This treatment had 6 replicates.
- 1 Transition and 34 days of desiccation: Sediments were dry during the preconditioning and transitions periods, and transitioned once to the field moisture level prior to respiration estimation. This treatment had 6 replicates.
- 2 Transitions and 4 days of desiccation: Sediments were held at field moisture levels for the preconditioning period and then for the first 7 days of the transitions period, then transitioned to a dried state for 4 days, and transitioned to field moisture conditions prior to respiration estimation. This treatment had 7 replicates.
- 3 Transitions and 31 days of desiccation: Sediments were dry during the preconditioning period and the first 4 days of the transitions period, then transitioned to field moisture levels for three days, then transitioned to 4 days in a dried state, and transitioned again to field moisture levels prior to respiration estimation. This treatment had 7 replicates.
- 4 Transitions and 8 days of desiccation: Sediments were held at field moisture levels for the preconditioning period and transitioned to a dried state for the first 4 days of the transitions period, then transitioned to field moisture levels for 3 days, then transitioned to 4 days in a dried state, and transitioned to field moisture levels prior to respiration estimation. This treatment had 7 replicates.
- 5 Transitions and 27 days of desiccation: Sediments were dry during the preconditioning period and then transitioned to field moisture levels for the first 3 days of the transitions period, transitioned to a dried state for 2 days, transitioned to field moisture levels for 2 days, transitioned to a dried state for 2 days, and transitioned to field moisture levels for 3 days prior to respiration estimation. This treatment had 7 replicates.

**Figure 2.**
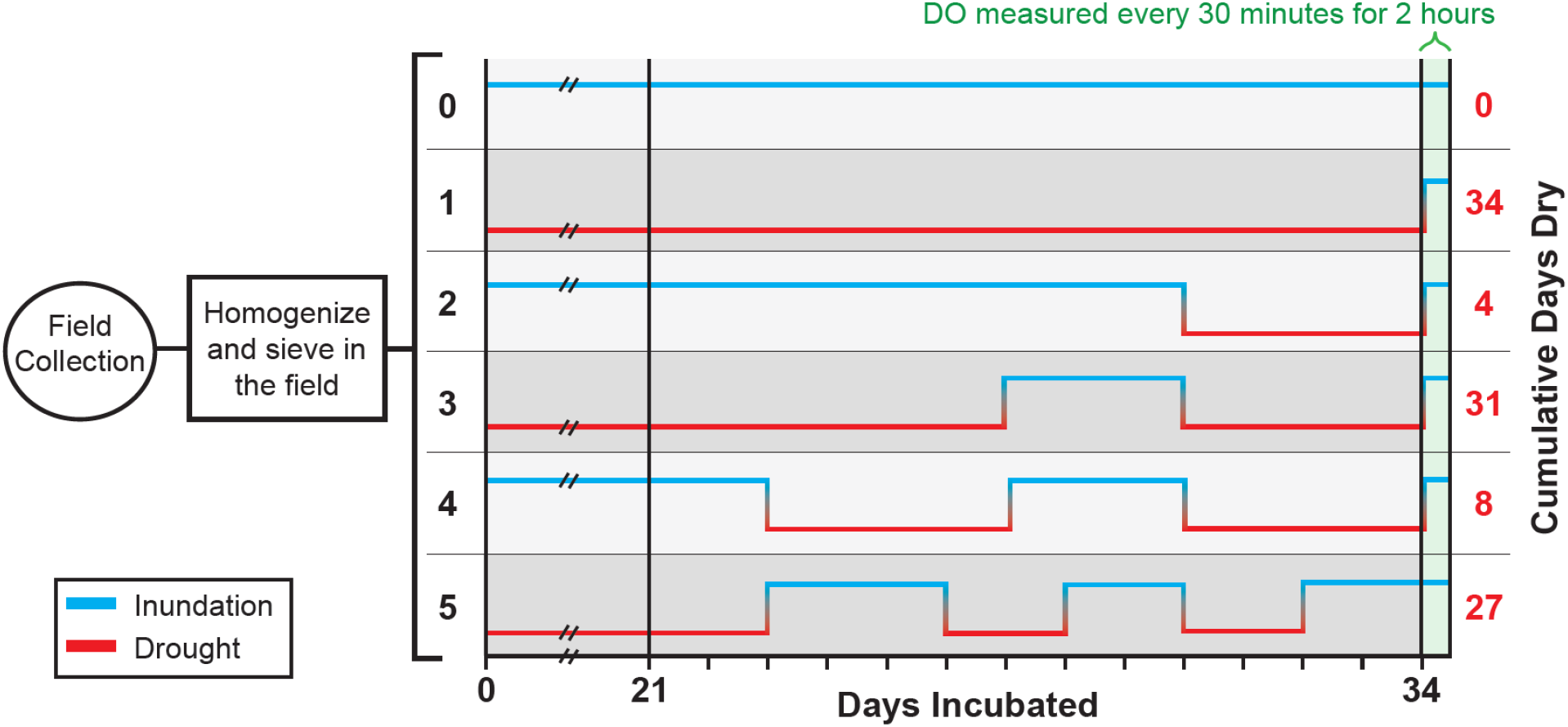
Experimental design of batch reactor incubations subjected to six treatment regimes of inundated (blue line) and dry (red line) conditions. Black values on the left indicate the number of inundated/dry transitions, including the final inundation that all treatments experienced immediately prior to the measurement of respiration. Red values on the right indicate the number of cumulative dry days (e.g., treatments with 1 or 3 transitions experienced 34 or 31 cumulative dry days, respectively. Transitions between inundated and dry conditions started on day 24. All treatments were held at either an inundated or dry state prior to day 24.

To avoid modifying electrical conductivity across experimental treatments, sterile deionized water was added to reactors to achieve/maintain field moisture levels according to the defined wet/dry regimes detailed above. For reactors with sediments that were below field moisture levels, deionized water was added to achieve field moisture levels prior to respiration rate estimation. Changes in the total mass of reactors and volumes of water added during the course of the experiment are provided in Data file S1.

### Respiration rate measurements

Laboratory incubations were performed in batch reactors to quantify dissolved oxygen consumption rates. Borosilicate glass vials (20 ml) (I-Chem™ Clear VOA Glass Vials, Thermo-Fisher, Waltham, MA) served as incubator reactors. Factory calibrated oxygen sensor spots (Part# 200001875, diameter = 0.5 cm, detection limit 15 ppb, 0 – 100 % oxygen; PreSens GmbH, Regensburg, Germany) were adhered to the inner vials of the reactor prior to sediment addition. Detailed description of sensor adhesion and non-destructive measurements of DO consumption using these sensors is provided in Garayburo-Caruso et al. (*49*). Vials were grouped into six treatment regimes (explained in the previous section) representing inundation-drought transitions.

Sample processing and incubations were performed in a laboratory at 21±1 °C. The reactors were monitored for 2 hours, with measurement of dissolved oxygen (DO) concentration (μmol L^−1^) every 30 min. DO concentration in each bioreactor was measured with an oxygen optical meter (Fibox 3; PreSens GmbH) connected to a 2 mm polymer optical fiber lined up to sense the sensor dot every thirty minutes. Respiration rates (μmol L^−1^ h^−1^) were estimated as the slope of the linear regression between DO concentration and incubation time for each sample. Some non-linearity was observed in the relationship between DO concentration and time such that only the first 4 data points--time zero to 2 hours--were used to fit a linear function. The slope of the linear function was taken as an estimate of respiration rate.

### Microbial Analysis

Post-incubation, the sediment slurry was transferred to centrifuge tubes (Item#28-108 Genesee Scientific) and centrifuged for 5 min at 3200 rcf and 20°C. The supernatant was removed and reserved for biogeochemistry analyses and sediment aliquots for DNA and RNA extraction were flash-frozen in liquid N_2_ and stored at −80 °C. The extraction, purification, and sequencing of sediment microbial gDNA were performed according to published protocol (*29*). The extraction of RNA was performed using the Qiagen PowerSoil RNA extraction kit (Qiagen, Germantown, MD). RNA was treated with DNase and quantified with a Qubit RNA kit (Thermo Fisher, Waltham, MA). An aliquot of the RNA extraction was used to generate cDNA using SuperScript™ IV First-Strand Synthesis System (Thermo Fisher Scientific, Waltham, MA). The 16S rRNA gene sequencing--for both gDNA and cDNA--followed the established protocol by The Earth Microbiome Project (*94*). Sequence pre-processing, operational taxonomic unit (OTU) assignment, and phylogenetic tree building were performed using an in-house pipeline, HUNDO (*69*). Sequences were deposited at NCBI’s Sequence Read Archive PRJNA641165. The final sample count of gDNA and cDNA, respectively, for each treatment regime, after dropping samples following quality filtering and rarefaction, was 5 and 5 (0 transition), 4 and 3 (1 transition), 5 and 4 (2 transitions), 7 and 6 (3 transitions), 7 and 6 (4 transitions), and 7 and 5 (5 transitions). Rarefaction levels are provided below in the Statistics section.

### Biogeochemistry

Reserved supernatant was filtered through a 0.22 μm polyethersulfone membrane filter (Millipore Sterivex) and an aliquot was immediately removed for non-purgeable organic carbon (NPOC) and the remainder was stored at −20C until further OM high resolution analysis was conducted (see below). NPOC was determined by acidifying an aliquot of sample with 15% by volume of 2N ultra-pure HCL (Optima grade, Fisher#A466-500). The acidified sample was sparged with carrier gas (zero air, Oxarc# X32070) for 5 minutes to remove the inorganic carbon component. The sparged sample was then injected into the TOC-L furnace of the Shimadzu combustion carbon analyzer TOC-L CSH/CSN E100V with ASI-L auto sampler at 680°C using 150 uL injection volumes. The best 4 out of 5 injections replicates were averaged to get the final result. The NPOC organic carbon standard was made from potassium hydrogen phthalate solid (Nacalia Tesque, lot M7M4380). The calibration range was 0 to 70 ppm NPOC as carbon.

Fourier Transform Ion Cyclotron Resonance Mass Spectrometry (FTICR-MS) of post-incubation sediment slurry was conducted as per Danczak et al. (*70*). Sample processing, injection, and data acquisition, processing and analysis was performed as per scripts provided in Danczak et al. (*70*), with ‘*Start tolerance’* in Formularity changed to 8. Ten samples were dropped due to poor calibration, resulting in 5 replicates for 0 transition, 4 replicates for 1 transition, 5 replicates for 2 transitions, 6 replicates for 3 transitions, and 5 replicates each for 4 and 5 transition regimes.

From the FTICR-MS data, as in previous work (*32, 34, 47, 49*), we followed LaRowe and Van Cappellen (*71*) to calculate the Gibbs free energy for the half reaction of organic carbon oxidation under standard conditions (ΔG^0^_Cox_). This calculation is based on elemental stoichiometries associated with molecular formulae assigned to individual molecules observed in the FTICR-MS data. The formulae assignments are part of the processing scripts described in (*70*). As in previous work (*32, 34, 47, 49*), we followed LaRowe and Van Cappellen (*71*), we interpret larger values of ΔG^0^_Cox_to indicate OM that is thermodynamically less favorable for oxidation by microbes. That is, larger values of ΔG^0^_Cox_ indicate OM that provides less net energy to a microbial cell per oxidation event, assuming all else is equal. Given the large numbers of assigned formulae within each sample, this resulted in thousands of ΔG^0^_Cox_ estimates within each sample, from which we estimated mean ΔG^0^_Cox_ for each sample.

## Statistics

### Estimating influences of community assembly processes

The relative influences of community assembly processes influencing microbial community membership are emergent properties that cannot be calculated/inferred directly from knowledge of membership. To evaluate assembly processes as a link between membership and microbial processes (reference framework graphic) it is necessary to quantitatively estimate the relative influences of these processes. To do so we use a well-established null modeling framework based on phylogenetic relationships among microbial taxa (*22, 24, 25, 27*). We refer the reader to these previous studies for details. In brief, randomizations were used to generate estimates of phylogenetic associations among microbial taxa for scenarios in which microbial communities were stochastically assembled. These stochastic (i.e., null) expectations were compared quantitatively to observed phylogenetic associations to estimate the *β*-Nearest Taxon Index (*β*NTI) (*22*). We used cDNA sequences rarefied to 27227 and gDNA sequences rarefied to 15106 sequences per sample to determine putatively active community and whole community *β*NTI values, respectively. Samples falling below these sequence counts were removed as indicated above in subsection *Microbial Analysis.* A *β*NTI value of 0 indicates no deviation between the stochastic expectation and the observed phylogenetic associations, thereby indicating the dominance of stochastic assembly processes. As *β*NTI deviates further from 0, there is an increasing influence of deterministic assembly processes that drive community membership away from the stochastic expectation. *β*NTI values below −2 or above +2 indicate statistical significance, with negative and positive values indicating less than or more than expected shifts in membership. *β*NTI is a pairwise metric measured between any two communities/samples, such that shifts in membership are related to changes between the pair of communities being evaluated. We used *β*NTI to study all pairwise community comparisons within each experimental treatment. Each community from a given reactor is therefore associated with multiple *β*NTI values due to being compared to communities associated with other replicate reactors. In turn, the average *β*NTI was calculated for each reactor. As in Stegen et al. (*25*), this provides a community-specific value for *β*NTI and thus an estimate of the relative influences of stochastic and deterministic processes causing deviations between a given community and all other communities within the same experimental conditions (*32*). That is, the larger the absolute value of *β*NTI for a given community, the stronger the influence of deterministic assembly processes acting on that community (*25*). In turn, these community-specific estimates were related to reactor-specific measurements. For example, respiration rates were regressed against *β*NTI to evaluate the link between emergent properties (i.e., ecological assembly) and microbial processes (i.e., respiration rates).

### Evaluating relationships between microbial characteristics and environment

Respiration rate distributions, absolute *β*NTI values, and ΔG^0^_Cox_ were summarized as box plots. Pairwise Mann-Whitney test was performed to evaluate statistical differences between reactor-specific measurements (*e.g.*, respiration rates and thermodynamic properties) and treatment groups (cumulative dry and inundated days). Continuous bivariate relationships were evaluated with ordinary least squares regression. Prior to regression analyses, respiration rates were log-transformed due to non-linearities resulting from respiration being constrained to be at or above zero. Prior to log-transformation, half the smallest non-zero rate was added to each rate to enable inclusion of rate estimates with a value of zero.

### Control point influence calculation

To characterize respiration rate distributions we used the control point influence (CPI) metric. CPI was recently developed (*53*) and is defined as the fraction of cumulative function (R_tot_; e.g., total respiration rate) within a defined system that is contributed by individual rates that are above the system’s median rate (R_med_). To define cumulative function one must first define the system being evaluated. In our study, all replicate batch reactors within a given experimental treatment were conceptualized as a representative set of samples from a larger system experiencing the experimental conditions. R_tot_ for each treatment was therefore estimated as the sum of respiration rates across a treatment’s replicate reactors. CPI was estimated as the sum of respiration rates that fell above the median rate for a given treatment (R_above_) divided by R_tot_ for that treatment. That is, 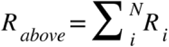, where R_i_ are respiration rates from individual reactors that fell above R_med_, and 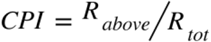.

An important feature of CPI is that it makes no assumptions of distribution normality, and can be estimated for rate distributions of any form (e.g., unimodal, multimodal, Gaussian, skewed, etc.). In most cases, CPI is constrained to have a minimum value of 0.5 (for a perfectly normal distribution with no outliers) and asymptotically approach 1 as a maximum value (e.g., for heavily skewed distributions with a small number of very high rates). CPI therefore quantitatively estimates the biogeochemical contribution of places in space or points in time that have elevated biogeochemical rates.

## Results

To link microbial membership to emergent microbial community properties (Fig. 1 arrow 1) we used null modeling to estimate the contributions of stochastic and deterministic community assembly. Results from the null models indicate a relatively balanced mixture of stochasticity and determinism for both the whole community (gDNA-based) and putatively active community (rRNA-based) (Fig. S2). More specifically, stochasticity and determinism each governed 50% of turnover in microbial membership for the whole community and 33% and 67%, respectively, for the putatively active community. The relative contributions of the two deterministic components (homogeneous and variable selection) were strongly imbalanced. Homogeneous selection was responsible for 94% and 91% of the deterministic component for the whole and putatively active communities, respectively. The contributions of homogeneous and variable selection to the deterministic component must sum to 1, such that the variable selection was responsible for 6% and 9% of the deterministic component for the whole and putatively active communities, respectively.

As shown in Figure 1 (arrow 2), we hypothesized a link between microbial community properties and microbial processes realized as a relationship between the strength of determinism and respiration rates. Such a relationship was not observed for the whole community (Fig. 3a), but we did observe a non-linear decreasing relationship between respiration rates and the absolute value of βNTI for the putatively active community (Fig. 3b). The direction of this relationship (negative) was opposite of that expected.

**Figure 3.**
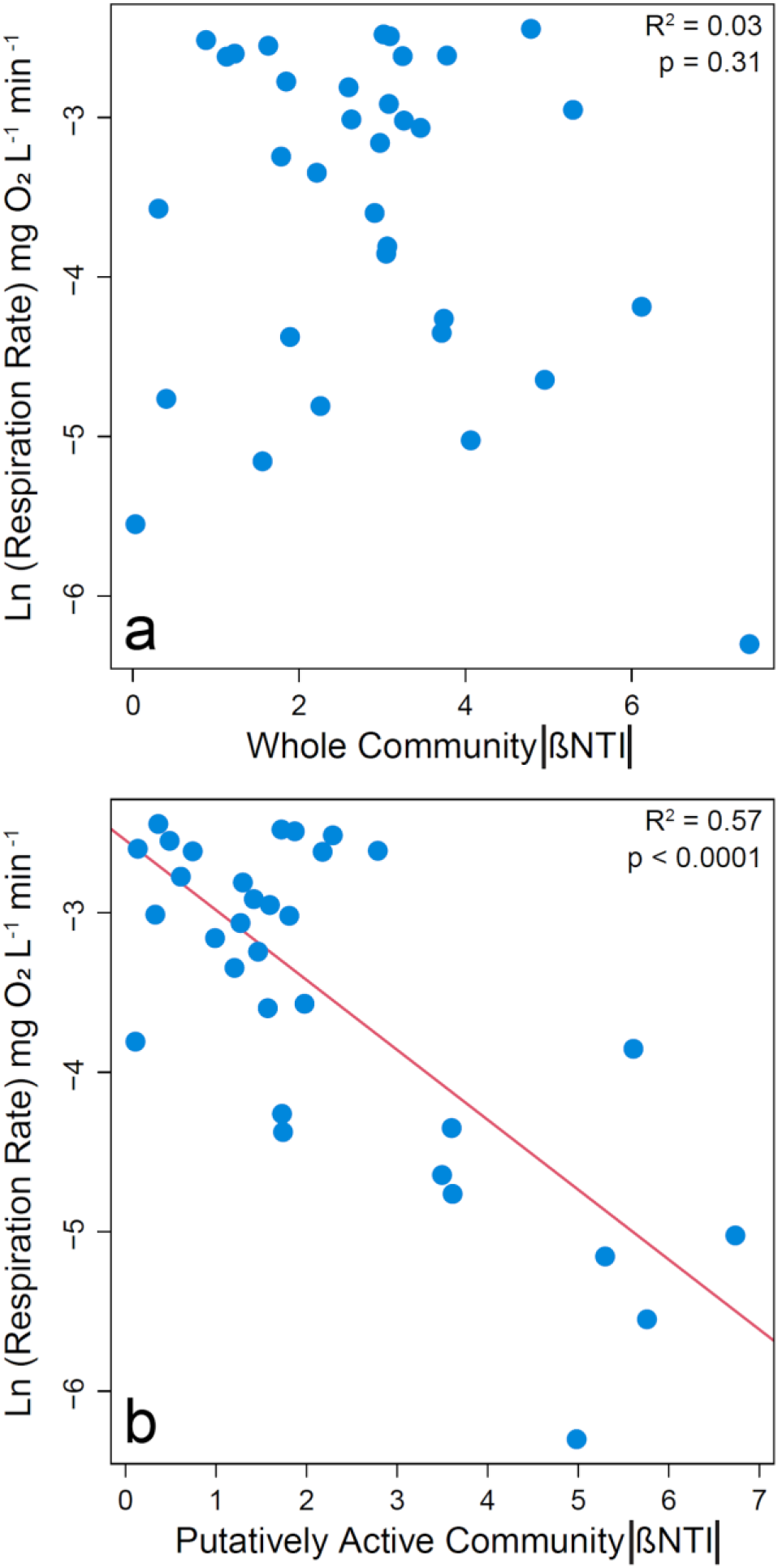
Natural log transformed respiration rates (i.e., O_2_ consumption) as a function of the absolute value of βNTI for (a) whole communities or (b) putatively active communities. Larger absolute values of βNTI indicate stronger influences of deterministic assembly. Nonlinearity was observed because the respiration rate has a lower limit of 0 such that its relationship with βNTI was fit as a negative exponential function. The significant regression model is shown as a red line, and statistics are provided on each panel.

In our conceptual framework there are multiple ways in which connections among the environment, microbial properties, and microbial processes may be realized, in part due to the environment having multiple components relevant to our study (Fig. 1 arrows 3,4). More specifically, the environment was characterized here in terms of both disturbance (number of dry days; imposed by the experimental manipulation) and OM thermodynamics (ΔG^0^_Cox_; this is an emergent aspect of the environment).

Disturbance influenced both microbial properties and processes. These influences appeared to be non-linear with experimental treatments associated with the two largest number of dry days (31 and 34) causing decreases in respiration rates (Fig. 4a) and stronger influences of deterministic homogeneous selection for the putatively active community (Fig. 4b). Disturbance had no clear influence on community assembly for the whole community (Figs. S3, S4a). Given the apparent binary nature of these results, we evaluated statistical significance by combining respiration rate data from treatments with 0-27 cumulative dry days and separately combining data from treatments with 31 or 34 cumulative dry days (Fig. S5). Respiration rates were significantly depressed in the treatments associated with 31 or 34 cumulative dry days (W = 5, p < 0.001). The βNTI data are non-independent due to being based on all pairwise comparisons within a treatment. Standard statistical tools are therefore not applicable for assigning statistical significance when comparing βNTI distributions. However, as shown in Figures 4b and S4b, there is an obvious shift to lower βNTI values for the putatively active community in the treatments with 31 or 34 cumulative dry days.

**Figure 4.**
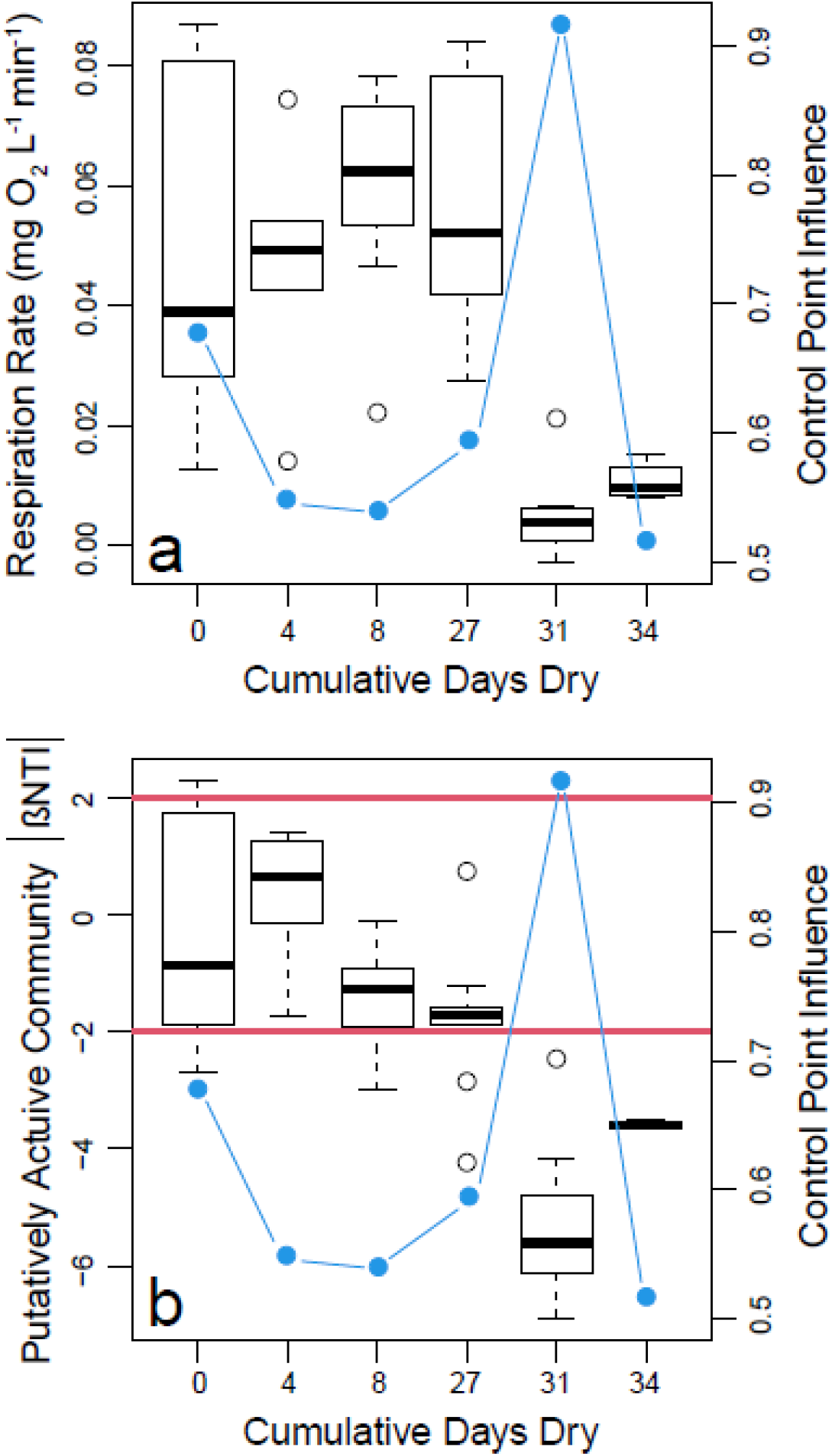
Boxplot representations of respiration rate (a) and putatively active community βNTI (b) distributions as a function of the cumulative number of days reactors were in a dried state. Each value along the horizontal axis represents a different experimental treatment. On both panels the right hand axis provides estimates of control point influence (blue circles and lines) across the treatments. Horizontal red lines in (b) indicate significance thresholds; values below −2 indicate deterministic homogenous selection, values above +2 indicate deterministic variable selection, and values between −2 and +2 indicate stochastic assembly.

The other aspect of the environment examined here (i.e., OM thermodynamics) also had significant relationships with both microbial processes (Fig. 1, arrow 3) and properties (Fig. 1, arrow 4). More specifically, respiration rates decreased significantly as a negative exponential function of increasing ΔG^0^_Cox_ (R^2^ = 0.26; p = 0.01; Fig. 5a). This indicates a decrease in respiration rate as OM thermodynamic properties shifted towards lower favorability for oxidation (i.e., larger values of ΔG^0^_Cox_). Similarly, we found that the strength of deterministic assembly associated with the putatively active community increased linearly with ΔG^0^_Cox_ (R^2^ = 0.46; p < 0.0001; Fig. 5b). The strength of deterministic assembly associated with the whole community was unrelated to ΔG^0^_Cox_ (p = 0.64) (Fig. S6).

**Figure 5.**
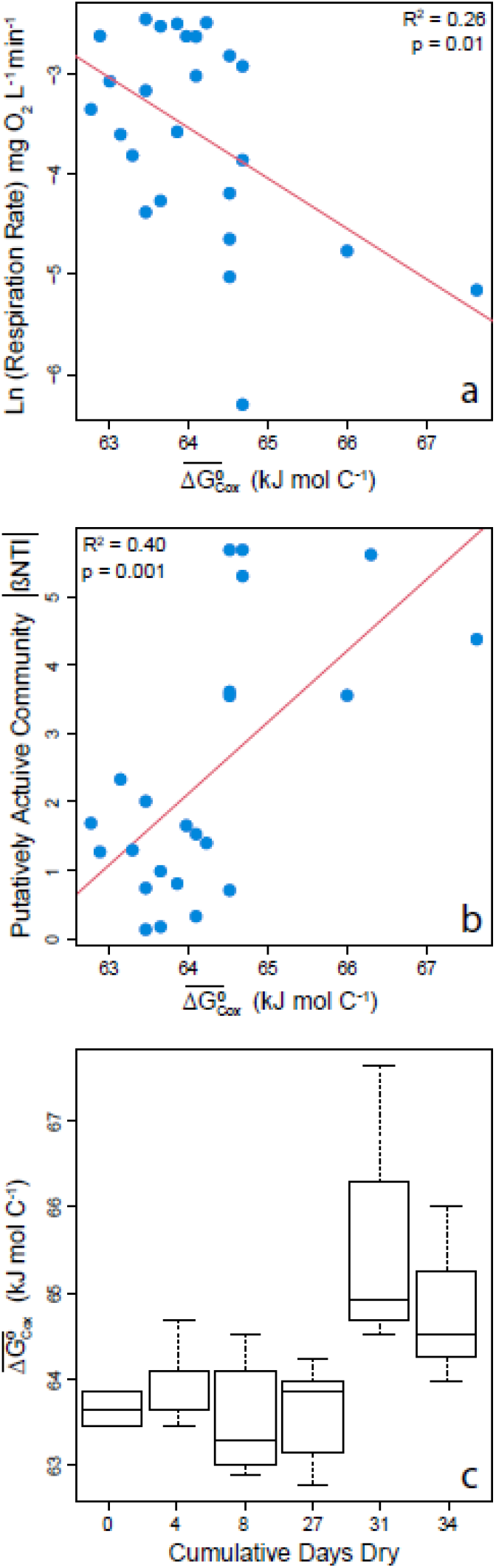
Microbial processes and properties as a function of OM thermodynamics, and impacts of disturbance on OM thermodynamics. (a) Respiration rates (natural log transformed) decreased with decreasing favorability for oxidation (larger values of ΔG^0^_Cox_). (b) The strength of deterministic selection measured as the absolute value of βNTI increased with decreasing favorability for oxidation. Regression models are shown as red lines and statistics are provided on each panel. (c) Boxplot representations of the distributions of OM thermodynamics across experimental treatments. Significant increases were observed for treatments with 31 or 34 cumulative dry days. See text and Figure S7 for a description of statistics.

The conceptual model described in Figure 1 focuses primarily on connections among environmental and/or microbial attributes, but there are potentially important relationships within attribute categories. In particular, within the environmental category there is the potential for an influence of disturbance on OM thermodynamics. Such an effect was found for OM thermodynamics as measured by ΔG^0^_Cox_ (Fig. 5c). Using the same approach as for analyses described above, we combined data for treatments with 0-27 cumulative dry days and compared that distribution to data combined across treatments with 31 or 34 cumulative dry days. A Mann-Whitney test comparing these distributions confirmed a significant change in the ΔG^0^_Cox_ distribution (W = 189, p = < 0.001)(Fig. S7).

The last component of the conceptual model considered here is the connection between microbial processes occurring in a given location and cumulative system function that aggregates across locations (Fig. 1, arrow 5). It is at the system level that the influence of biogeochemical hot spots (or hot moments) can be evaluated. We conceptualized an aggregate system as the collection of replicate batch reactors within a given experimental treatment. Based on this definition, we estimated control point influence (CPI) as a measurement for the influence of biogeochemical hot spots. We observed a large amount of variation in CPI across experimental treatments, but there was no clear, direct influence of the treatments on CPI (Fig. 4). The largest value of CPI observed (> 0.9) was associated with the treatment that imposed 31 cumulative dry days. This treatment also had the lowest median respiration rates across all treatments (Fig. 4).

## Discussion

Mechanistic evaluation of microbe-environment interactions is fundamental to understanding microbe-mediated ecosystem function. Inspired by a microbe-environment-ecosystem framework proposed by Hall et al. (*6*) we proposed and evaluated a modified framework linking microbial characteristics (membership, emergent properties, processes), the environment (disturbance, OM thermodynamics), and cumulative ecosystem function (CPI) of hyporheic zone sediments. Our results provide clear support for the overall conceptual framework and further point to an iterative loop among OM thermodynamics, respiration rates, and microbial community assembly that can be initiated by externally-imposed disturbance. Furthermore, our results indicate that the iterative thermodynamics-assembly-respiration loop may be initiated through threshold-like impacts of disturbance that were observed only after 31 or more cumulative days of desiccation.

We first evaluated emergent community properties as a function of microbial membership by studying the relative influences of stochasticity and determinism over community assembly. Taking this approach, we found fully balanced stochastic-deterministic influences over the whole community, in which each contributed to 50% of the variation in community composition. The relative influences of stochasticity and determinism have been quantified for many microbial systems and the estimates are highly variable (*72, 73*). In addition, within the deterministic component of assembly, homogeneous selection had a far greater influence than variable selection. Previous work has also observed a broad range of contributions from homogeneous and variable selection (*63, 74–77*). As such, the assembly-associated outcomes observed here for the whole community are not unexpected relative to previous work. Very few studies, however, have examined the relative influences of different assembly components over putatively active microbial communities.

For the putatively active communities we found that across all treatments both stochasticity and determinism were important, though deterministic assembly had greater influence. This deviates quantitatively from the whole community in which the influences of stochasticity and determinism were more balanced. Consistent with the whole community results, however, was the dominance of homogeneous selection within the deterministic component of assembly. The strong influence of homogeneous selection is likely due to selection-based constraints imposed by aspects of the experimental system that did not vary across treatments. For example, mineralogy is known to strongly influence microbial communities (*35, 78–82*) and was homogenized across the experimental batch reactors, thereby potentially imposing homogeneous selection on both the whole and putatively active communities.

Our study uniquely evaluates null-model outcome of putatively-active community assemblies in hyporheic zone sediments, where homogeneous selection was further enhanced by our experimentally imposed hydrologic disturbances. Increased homogeneous selection in response to disturbance is consistent with previous work in a soil system. In the soil system, disturbance led to an immediate increase in homogeneous selection for the putatively active community (*31*). The strong influence of homogeneous selection on the putatively active community is not always observed, however, suggesting it may be tied to acute disturbance. That is, Jia et al. (*42*) recently found that within a natural soil chronosequence, variable selection was stronger for putatively-active communities while homogeneous selection influenced the whole community assembly. Our results combined with these previous studies indicate that community assembly of putatively-active members may be more closely linked to short-term environmental change than assembly of the whole community.

In addition to being more sensitive to disturbance, we find that assembly of the putatively active community was more strongly tied to microbial processes (i.e., respiration rate), than was the whole community. A strong link between biogeochemical rates and the putatively active community is consistent with previous studies (*83, 84*). More specifically, we observed a negative relationship between respiration rate and absolute values of βNTI for the putatively active community, but no relationship for the whole community. The direction of this relationship is opposite to our hypothesis. While stronger selection should remove mal-adapted individuals, leading to higher biogeochemical rates (*23*), increased selection in our experiment was imposed by disturbance that appeared to directly suppress respiration rates due to desiccation (*85, 86*). The simultaneous suppression of respiration and imposition of stronger selection led to the negative relationship between respiration and the strength of selection. The lack of such relationships when considering the whole community indicates that a greater focus on assembly dynamics of putatively active communities could reveal new insights into the multi-component linkages among microbes, the environment, and function.

Disturbance also impacted OM thermodynamics and respiration rates, potentially initiating an iterative loop among microbial assembly, microbial processes, and the abiotic environment. In this iterative loop the direction of causation between OM thermodynamics and microbial processes (Fig. 1, arrow 3) is not clear due to feedbacks, though we interpret a direction of causation from OM thermodynamics to microbial properties in terms of community assembly (Fig. 1 arrow 4). As such, there may be a loop between OM thermodynamics and microbial processes (i.e., respiration) embedded in a larger loop that also includes microbial properties (i.e., community assembly). Such feedbacks are inherent in complex systems and often lead to non-linear dynamics as observed here in terms of the threshold-like impact of desiccation on multiple system components (*87, 88*).

As key elements of the inferred system of feedbacks, the links among OM thermodynamic properties, respiration, and desiccation found here are consistent with recent work tied to the same field system. That is, Garayburu-Caruso et al. (*49*) also showed decreasing aerobic respiration with decreasing favorability for oxidation (i.e., larger values of ΔG^0^_Cox_) using sediments sourced ~2 years previously from the same field system. In addition, the impacts of desiccation found here are similar to Goldman et al. (*62*) after re-inundation. This impact of desiccation on respiration contrasts with the Birch effect (*89*) in soils whereby desiccation followed by re-wetting leads to enhanced respiration. The consistency across hyporheic zone studies and deviation from classical soil phenomena points to potential consistency in governing processes within the hyporheic zone that deviate from processes operating in soil systems. Further evaluation is needed across additional hyporheic zone systems to rigorously evaluate this inference, however.

In addition to linkages between the environment and microbial aspects of the system, our study revealed connections within the environmental components of the conceptual framework. That is, greater cumulative desiccation caused an increase in ΔG^0^_Cox_, indicating a significant change in OM thermodynamics (Fig. 7). While our data cannot pinpoint governing mechanism(s), we hypothesize that the ΔG^0^_Cox_ response may have been tied to increased ion concentration following desiccation. For example, OM chemistry may have been altered due to changes in abiotic sorption, limitations of microbially accessible C due to water potential constraints, and/or osmolyte production and formation of extracellular polymeric substances (*90–92*).

Irrespective of mechanisms, the shift in OM thermodynamics in response to desiccation was associated with a decline in respiration. We infer a causal connection between OM thermodynamics and respiration, potentially triggered by desiccation-driven shifts in OM chemistry and/or microbial physiology. This causal connection is supported by recent work (*49*) and the observation of a continuous function between ΔG^0^_Cox_ and respiration rate that transcended experimental treatments. Desiccation therefore likely influences and may even initiate an iterative loop among OM thermodynamics, microbial assembly, and biogeochemistry that underlies cumulative system function.

Cumulative system function can often be driven by ecosystem control points (*52*), but we observed relatively little indication of such behavior. That is, estimates of control point influence (CPI) were relatively low across most treatments. CPI is theoretically constrained to range from 0.5-1, with lower values indicating smaller influences of control points. In our study, all but one treatment had CPI between ~0.5 and 0.7. The associated distributions of respiration rates did not contain obvious outliers such that we interpret CPI values in the 0.5-0.7 range to be relatively low and not strongly influenced by control points or biogeochemical hot spots/moments (*51*). The treatment with 31 cumulative days of desiccation diverged from the rest in having a CPI value of ~0.9. This large CPI was due to a single outlier (Fig. 5a) such that most of the cumulative respiration across reactors was contributed by that single reactor. We interpret that single reactor as a biogeochemical hot spot or control point within that experimental treatment. It is unclear, however, what led to such behavior as disturbance did not have any systematic influence on CPI.

A strength of CPI as a metric is that it allows for direct quantitative comparisons across studies, systems, and scales. Ours is the first study to estimate CPI, however, such that we cannot yet make comparisons to previous work. Through future comparisons it will be possible to evaluate the strengths, weaknesses, and behavior of CPI. We expect that some patterns may emerge such as CPI having a greater likelihood to reach very high values (near 1) in systems with relatively low rates on average. In these conditions, even a modest quantitative increase in biogeochemical rates can lead to a large proportional change such that most cumulative function is from a single point in space and/or time, resulting in large CPI. We also expect that some biogeochemical processes will show greater variation in CPI than others, potentially due to variation in degree of functional redundancy (*93*). For example, processes such as respiration can be performed by numerous microbial taxa (i.e., there is high functional redundancy), while others are more constrained to a relatively small number of taxa (e.g., ammonia oxidation). We hypothesize that CPI may be lower on average and less variable across systems and scales for biogeochemical processes with greater functional redundancy. Additional work will be needed to test this hypothesis.

In this study we coupled intrinsic characteristics of natural hyporheic zone sediments with imposed constraints in the form of desiccation to evaluate an *a priori* conceptual framework modified from Hall et al. (*6*). Our results demonstrated strong and often non-linear connections among desiccation, OM thermodynamics, assembly of the putatively active microbial community, and respiration rates. Collating our results points to further modification of the framework into an *a posteriori* conceptual model containing nested feedback loops (**Fig. 6**). This conceptual model is consistent with the recently proposed unification of microbial ecology around the concepts of external forcing, internal dynamics, and historical contingencies (*46*). That is, we hypothesize that external forcing imposed by desiccation initiates multiple internal loops that drive biological and chemical dynamics that, in turn, underlie respiration responses to re-wetting that are contingent on desiccation history. The development of conceptual models such as this is key to incorporating additional mechanistic detail into predictive simulation models (e.g., reactive transport codes). We encourage further evaluation and improvement of both our *a priori* and *a posteriori* concepts across environmentally divergent conditions to generate knowledge that is transferable across systems.

**Figure 6.**
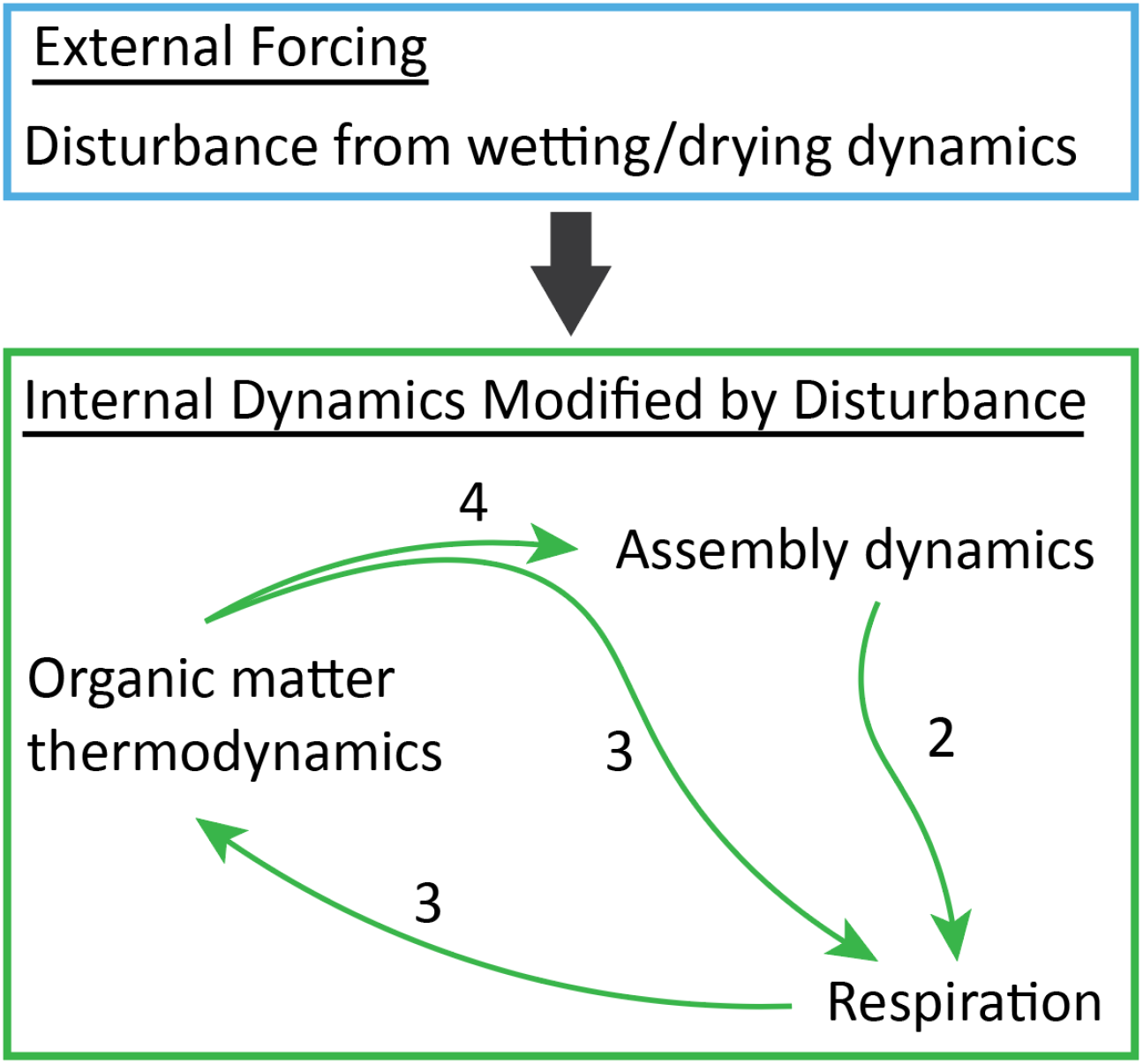
Integrated conceptual interpretation of results from this study. Collectively, our results indicate that the external forcing imposed by disturbance leads to feedback between assembly of the putatively active community and respiration rates that is modulated by coupled dynamics in organic matter thermodynamics. Relative to Fig. 1, here external and internal aspects of the environment are separated. The arrows within the internal dynamics component are analogous to arrows 2,3, and 4 in Figure 1. The arrow from external to internal is not considered in Figure 1, and represents the impact of external forcing on all aspects of the internal system. These impacts are both direct effects of disturbance and indirect effects mediated through the internal feedback that collectively lead to impacts of re-wetting that are contingent on desiccation history.

## Acknowledgements

We thank Amy Goldman and Nathan Johnson for developing graphics.

## Author contributions

JCS and SF conceptualized and designed the study; JT, JW, LR, RC, SF, and VAG-C performed the experiment; AS, JCS, RED, and SF analyzed data; AS, JCS, and SF drafted the manuscript and all authors contributed to further writing.

## Data and materials availability

Data are available on the ESS-DIVE archive at the following link (a data package will be published prior to the manuscript being published, and the associated link will be provided). Sequence data is available on NCBI’s Sequence Read Archive PRJNA641165.

## Supplementary Materials

**Supplementary Figure S1.**
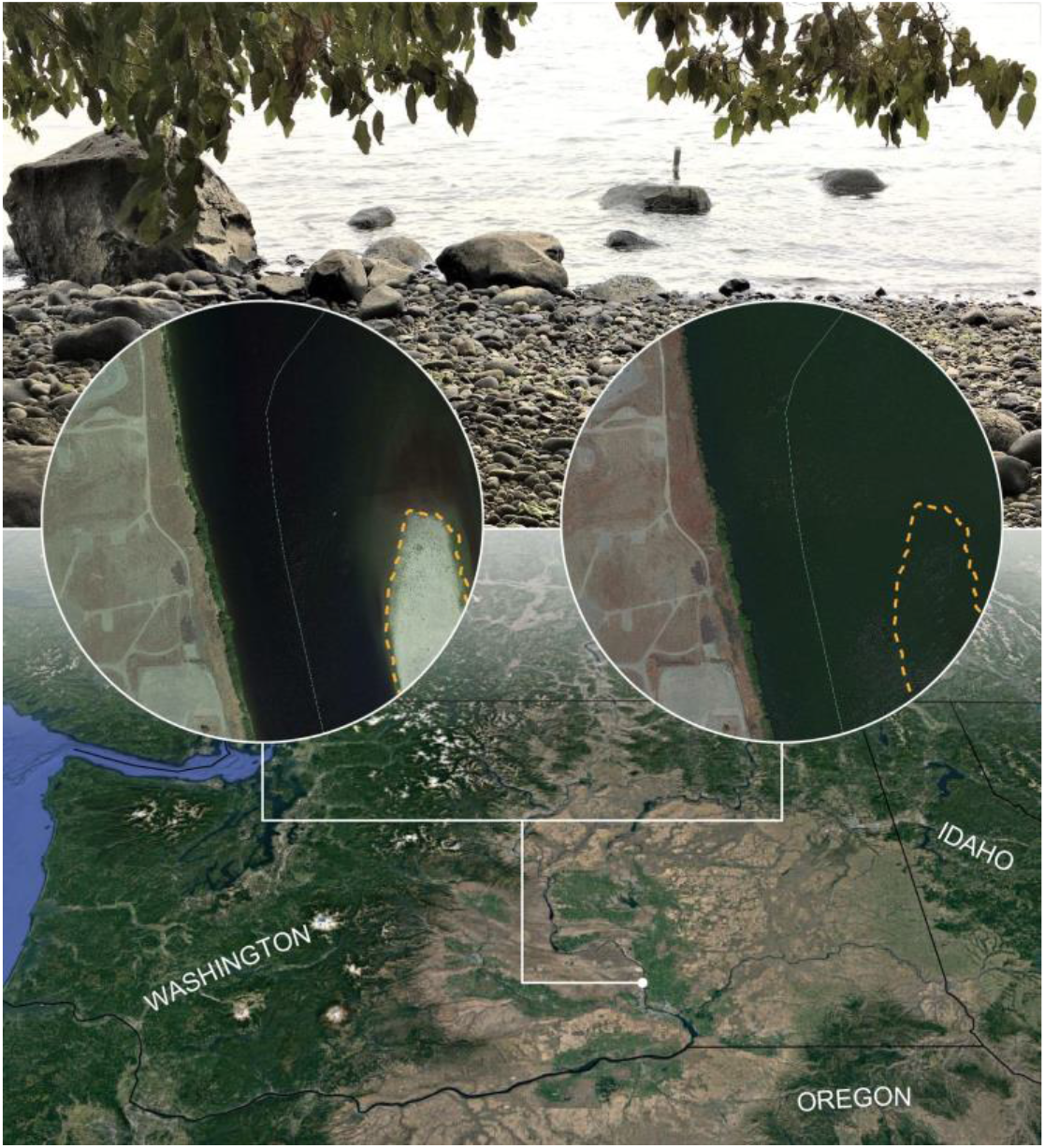
Field location from which sediments were collected. The site is along the Columbia River in southeast Washington State, as shown in the bottom portion of the image. The system is characterized by variable inundation of the river bed sediments and associated hyporheic zone. The middle images emphasize this variability in inundation. In those images the dashed yellow line is in the same location and outlines the shoreline of an island that is exposed during low water and inundated during high water. The photo at the top of the image is from the location of sediment collection. As can be seen, the system contains large cobbles and gravels, which are partially responsible for high hydrologic conductivity within the hyporheic zone, resulting in aerobic conditions during both inundated and non-inundated conditions.

**Supplementary Figure S2.**
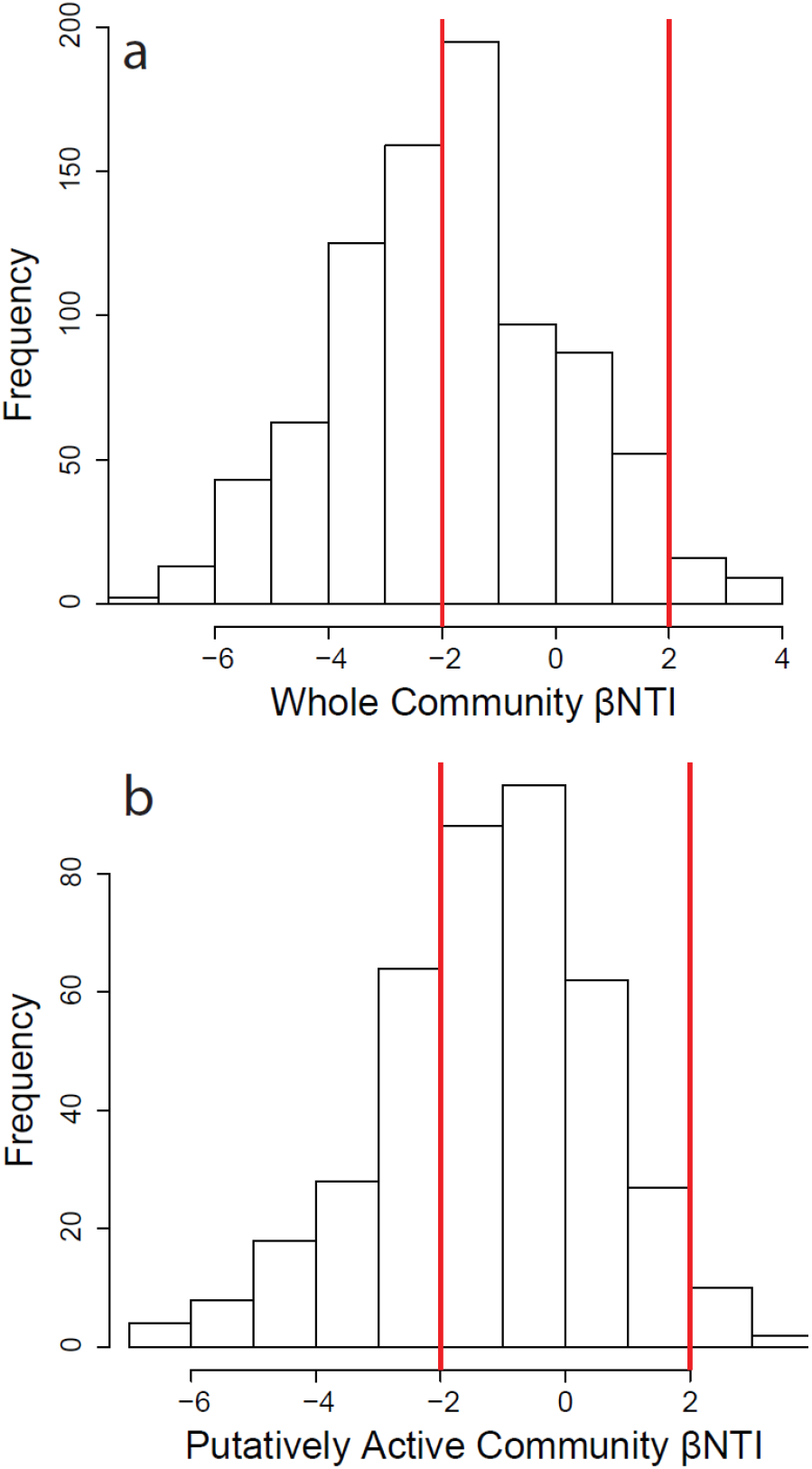
Histograms of βNTI associated with whole communities (a) and putatively active communities (b). Vertical red lines are the significant thresholds (−2 and +2). Values below −2 indicate deterministic homogenous selection, values above +2 indicate deterministic variable selection, and values between −2 and +2 indicate stochastic assembly.

**Supplementary Figure S3.**
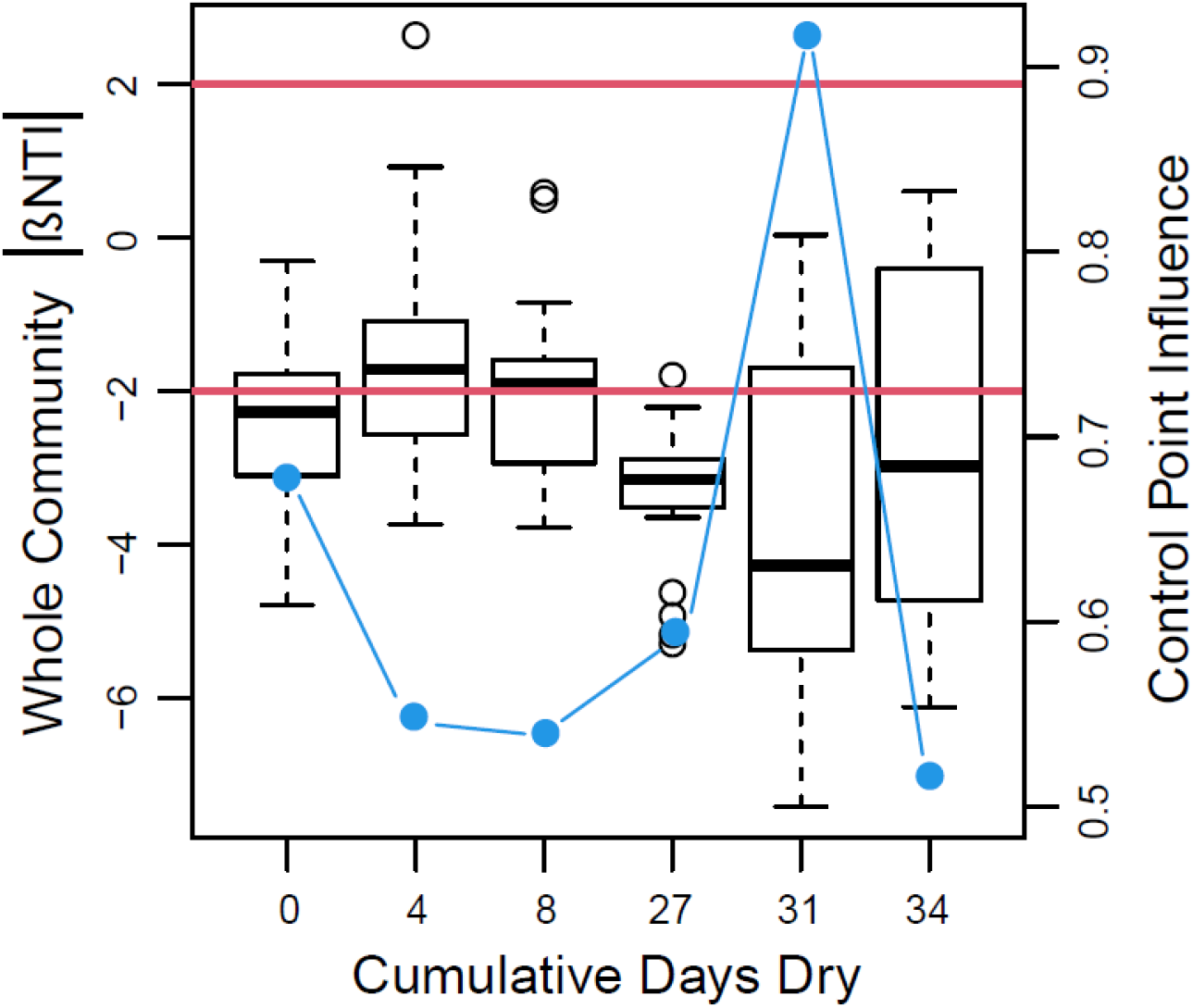
Boxplot representations of whole community βNTI distributions as a function of the cumulative number of days reactors were in a dried state. Each value along the horizontal axis represents a different experimental treatment. The right hand axis provides estimates of control point influence (blue circles and lines) across the treatments. Horizontal red lines indicate significance thresholds for βNTI; values below −2 indicate deterministic homogenous selection, values above +2 indicate deterministic variable selection, and values between −2 and +2 indicate stochastic assembly.

**Supplementary Figure S4.**
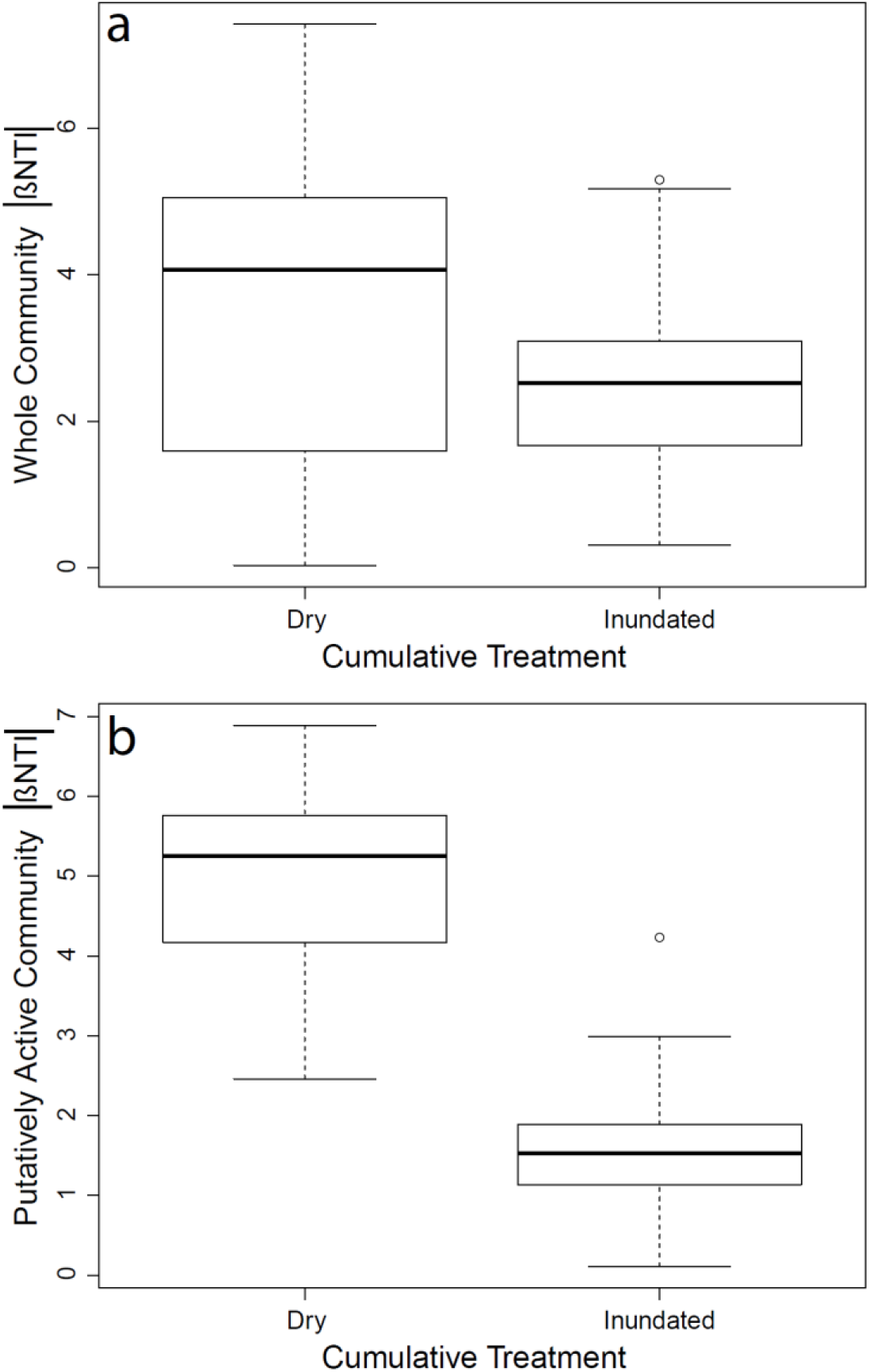
Boxplot representations of distributions of the absolute values of βNTI for the whole community (a) and the putatively active community (b) across two different categories of disturbance. On the horizontal axis ‘Dry’ indicates data combined from across treatments with 31 or 34 cumulative dry days, while ‘Inundated’ indicates data combined from across treatments with 0-27 cumulative dry days.

**Supplementary Figure S5.**
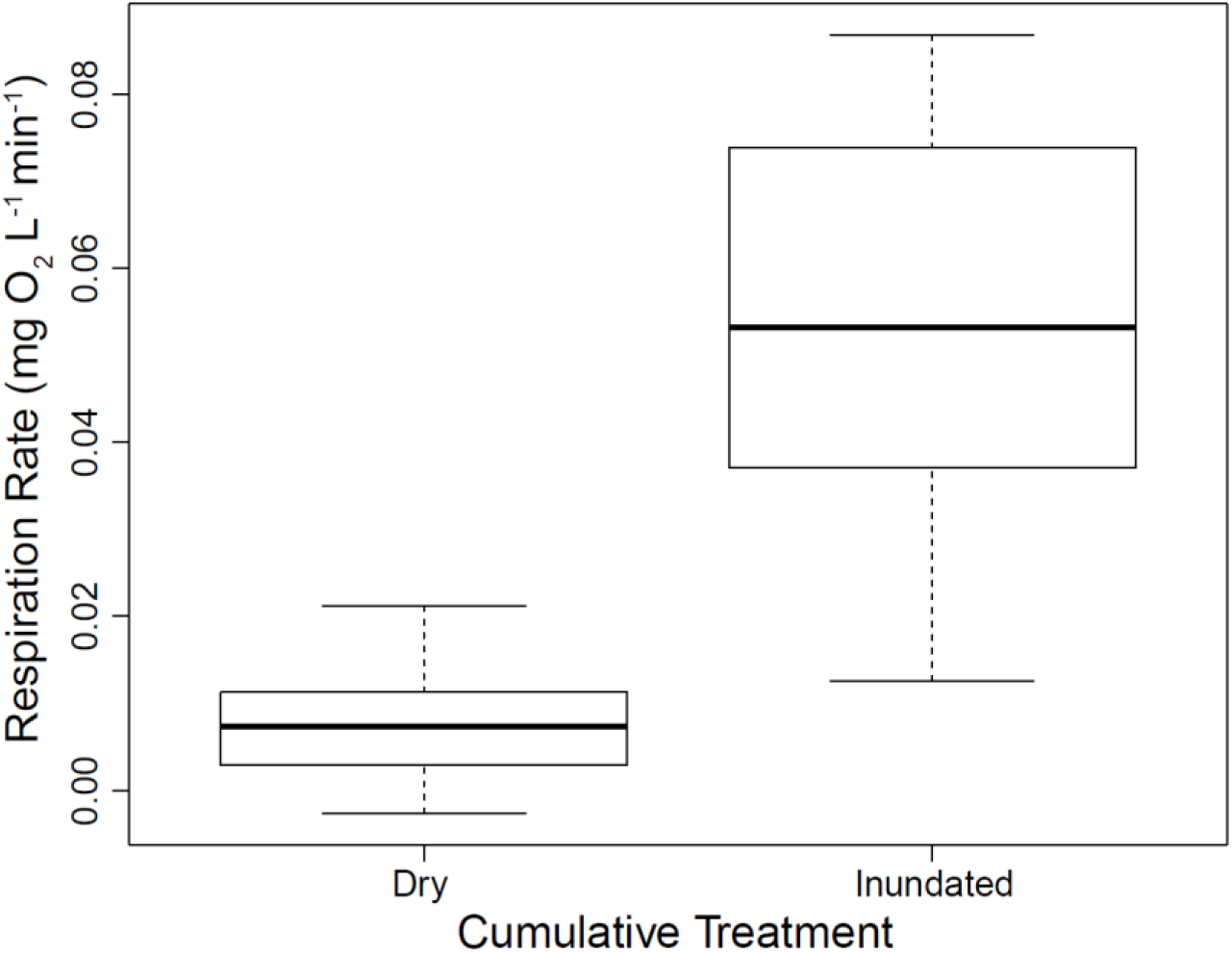
Boxplot representations of respiration rate distributions across two different categories of disturbance. On the horizontal axis ‘Dry’ indicates data combined from across treatments with 31 or 34 cumulative dry days, while ‘Inundated’ indicates data combined from across treatments with 0-27 cumulative dry days. The two distributions were significantly different per a pairwise Mann-Whitney test (W = 5, p < 0.001).

**Supplementary Figure S6.**
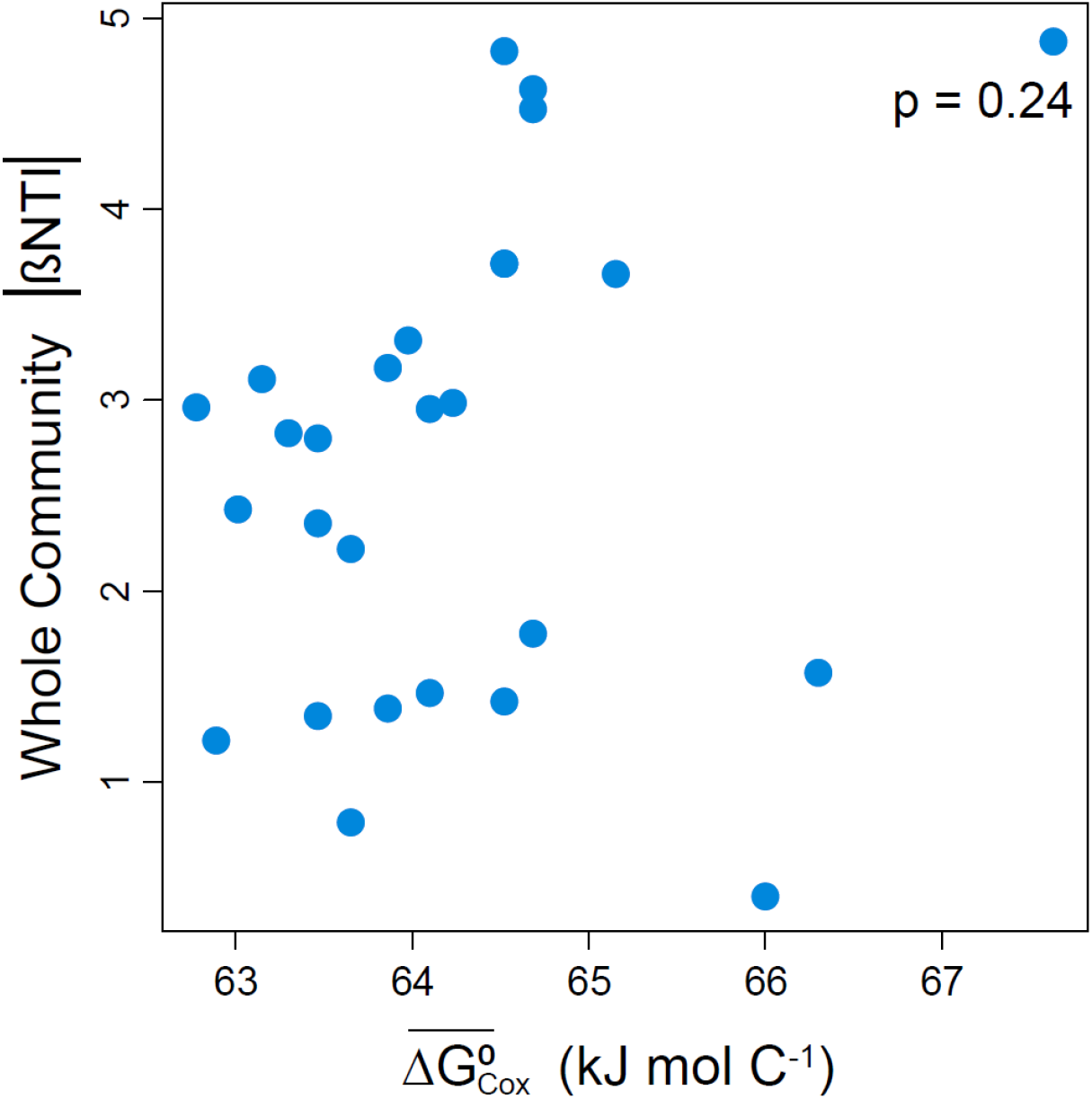
Community assembly as a function of organic matter thermodynamics. The strength of deterministic assembly associated with the whole community as measured by βNTI was not related to organic matter thermodynamics.

**Supplementary Figure S7.**
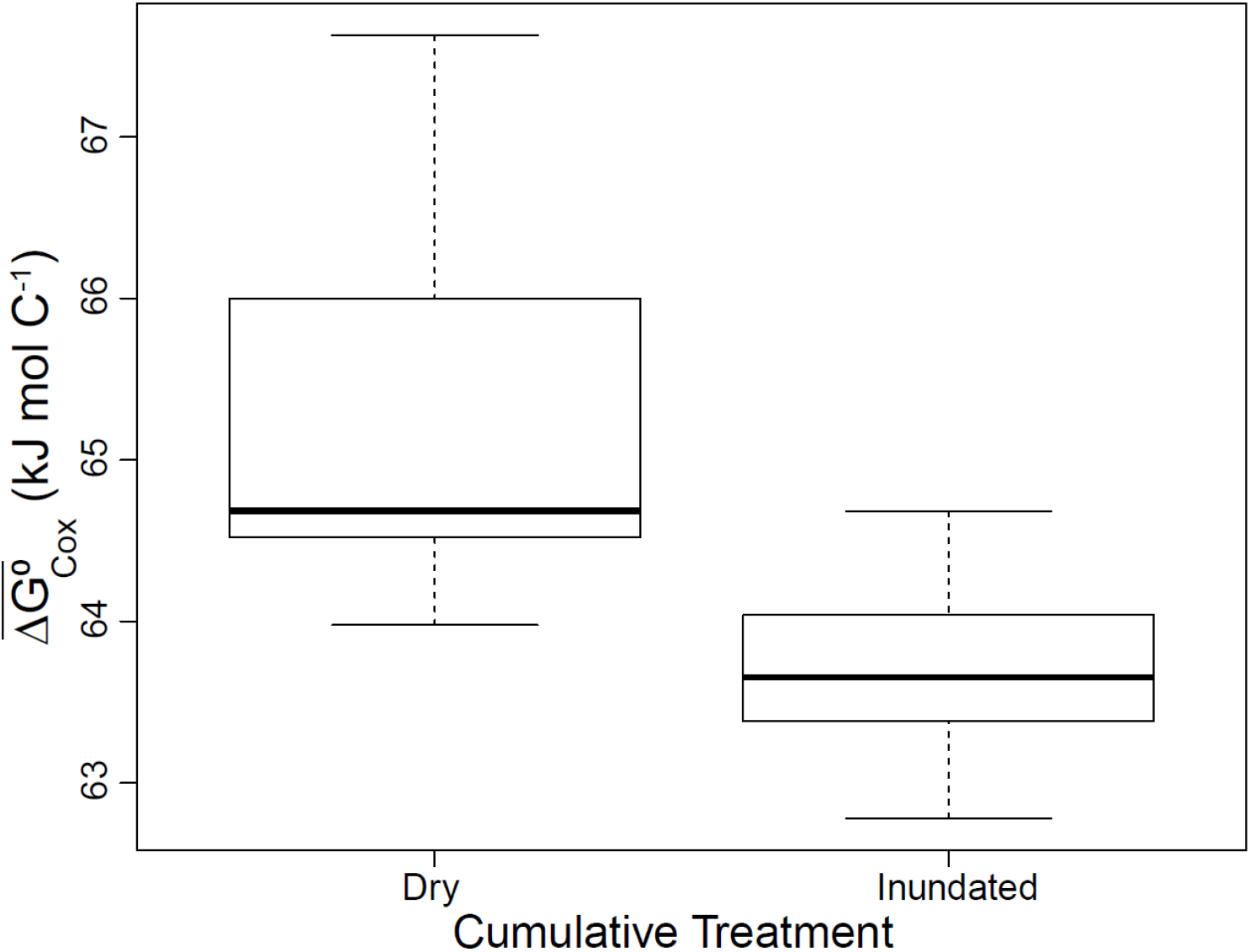
Boxplot representations of organic matter thermodynamics across two different categories of disturbance. On the horizontal axis, ‘Dry’ indicates data combined from across treatments with 31 or 34 cumulative dry days, while ‘Inundated’ indicates data combined from across treatments with 0-27 cumulative dry days. The two distributions were significantly different per a pairwise Mann-Whitney test (W = 189, p = < 0.001).

**Data file S1. Time series of moisture content within vial batch reactors.** Columns indicate the date (month/day/year), sample name that maps to the metadata file found on ESS-DIVE, the number of inundated/dessicated transitions (referred to as cycles for compatibility with R Scripts provided in ESS-DIVE data package), the estimated dry sediment mass in grams, and the mass of water in grams per gram of dry sediment.

## References

1. R. L. Bier, E. S. Bernhardt, C. M. Boot, E. B. Graham, E. K. Hall, J. T. Lennon, D. R. Nemergut, B. B. Osborne, C. Ruiz-González, J. P. Schimel, M. P. Waldrop, M. D. Wallenstein, Linking microbial community structure and microbial processes: an empirical and conceptual overview. FEMS Microbiol. Ecol. 91 (2015), doi:10.1093/femsec/fiv113.

2. A. A. Malik, J. B. H. Martiny, E. L. Brodie, A. C. Martiny, K. K. Treseder, S. D. Allison, Defining trait-based microbial strategies with consequences for soil carbon cycling under climate change. ISME J. 14, 1–9 (2020).

3. M. Manzella, R. Geiss, E. K. Hall, Evaluating the stoichiometric trait distributions of cultured bacterial populations and uncultured microbial communities. Environ. Microbiol. 21, 3613–3626 (2019).

4. M. D. Wallenstein, E. K. Hall, A trait-based framework for predicting when and where microbial adaptation to climate change will affect ecosystem functioning. Biogeochemistry. 109, 35–47 (2012).

5. W. R. Wieder, S. D. Allison, E. A. Davidson, K. Georgiou, O. Hararuk, Y. He, F. Hopkins, Y. Luo, M. J. Smith, B. Sulman, K. Todd-Brown, Y.-P. Wang, J. Xia, X. Xu, Explicitly representing soil microbial processes in Earth system models. Glob. Biogeochem. Cycles. 29, 1782–1800 (2015).

6. E. K. Hall, E. S. Bernhardt, R. L. Bier, M. A. Bradford, C. M. Boot, J. B. Cotner, P. A. del Giorgio, S. E. Evans, E. B. Graham, S. E. Jones, J. T. Lennon, K. J. Locey, D. Nemergut, B. B. Osborne, J. D. Rocca, J. P. Schimel, M. P. Waldrop, M. D. Wallenstein, Understanding how microbiomes influence the systems they inhabit. Nat. Microbiol. 3, 977–982 (2018).

7. J. A. Gilbert, J. K. Jansson, R. Knight, Earth Microbiome Project and Global Systems Biology. mSystems. 3 (2018), doi:10.1128/mSystems.00217-17.

8. J. Lloyd-Price, C. Arze, A. N. Ananthakrishnan, M. Schirmer, J. Avila-Pacheco, T. W. Poon, E. Andrews, N. J. Ajami, K. S. Bonham, C. J. Brislawn, D. Casero, H. Courtney, A. Gonzalez, T. G. Graeber, A. B. Hall, K. Lake, C. J. Landers, H. Mallick, D. R. Plichta, M. Prasad, G. Rahnavard, J. Sauk, D. Shungin, Y. Vázquez-Baeza, R. A. White, J. Braun, L. Denson, J. K. Jansson, R. Knight, S. Kugathasan, D. P. B. McGovern, J. F. Petrosino, T. S. Stappenbeck, H. S. Winter, C. B. Clish, E. A. Franzosa, H. Vlamakis, R. J. Xavier, C. Huttenhower, Multi-omics of the gut microbial ecosystem in inflammatory bowel diseases. Nature. 569, 655–662 (2019).

9. X. Fu, Y. Li, Y. Meng, Q. Yuan, Z. Zhang, D. Norbäck, Y. Deng, X. Zhang, Y. Sun, bioRxiv, in press, doi:10.1101/2020.01.05.893529.

10. S. König, A. Worrich, T. Banitz, F. Centler, H. Harms, M. Kästner, A. Miltner, L. Y. Wick, M. Thullner, K. Frank, Spatiotemporal disturbance characteristics determine functional stability and collapse risk of simulated microbial ecosystems. Sci. Rep. 8, 1–13 (2018).

11. S. Behrens, A. Kappler, M. Obst, Linking environmental processes to the in situ functioning of microorganisms by high-resolution secondary ion mass spectrometry (NanoSIMS) and scanning transmission X-ray microscopy (STXM). Environ. Microbiol. 14, 2851–2869 (2012).

12. S. Norland, K. M. Fagerbakke, M. Heldal, Light element analysis of individual bacteria by x-ray microanalysis. Appl. Environ. Microbiol. 61, 1357–1362 (1995).

13. L. R. Thompson, J. G. Sanders, D. McDonald, A. Amir, J. Ladau, K. J. Locey, R. J. Prill, A. Tripathi, S. M. Gibbons, G. Ackermann, J. A. Navas-Molina, S. Janssen, E. Kopylova, Y. Vázquez-Baeza, A. González, J. T. Morton, S. Mirarab, Z. Zech Xu, L. Jiang, M. F. Haroon, J. Kanbar, Q. Zhu, S. Jin Song, T. Kosciolek, N. A. Bokulich, J. Lefler, C. J. Brislawn, G. Humphrey, S. M. Owens, J. Hampton-Marcell, D. Berg-Lyons, V. McKenzie, N. Fierer, J. A. Fuhrman, A. Clauset, R. L. Stevens, A. Shade, K. S. Pollard, K. D. Goodwin, J. K. Jansson, J. A. Gilbert, R. Knight, A communal catalogue reveals Earth’s multiscale microbial diversity. Nature. 551, 457–463 (2017).

14. M. Wagner, Single-cell ecophysiology of microbes as revealed by Raman microspectroscopy or secondary ion mass spectrometry imaging. Annu. Rev. Microbiol. 63, 411–429 (2009).

15. R. P. Bartelme, J. M. Custer, C. L. Dupont, J. L. Espinoza, M. Torralba, B. Khalili, P. Carini, Influence of Substrate Concentration on the Culturability of Heterotrophic Soil Microbes Isolated by High-Throughput Dilution-to-Extinction Cultivation. mSphere. 5 (2020), doi:10.1128/mSphere.00024-20.

16. R. L. Barnard, C. A. Osborne, M. K. Firestone, Changing precipitation pattern alters soil microbial community response to wet-up under a Mediterranean-type climate. ISME J. 9, 946–957 (2015).

17. S. J. Blazewicz, R. L. Barnard, R. A. Daly, M. K. Firestone, Evaluating rRNA as an indicator of microbial activity in environmental communities: limitations and uses. ISME J. 7, 2061–2068 (2013).

18. D. C. Cardoso, A. Sandionigi, M. S. Cretoiu, M. Casiraghi, L. Stal, H. Bolhuis, Comparison of the active and resident community of a coastal microbial mat. Sci. Rep. 7, 1–10 (2017).

19. P. J. Kearns, J. H. Angell, E. M. Howard, L. A. Deegan, R. H. R. Stanley, J. L. Bowen, Nutrient enrichment induces dormancy and decreases diversity of active bacteria in salt marsh sediments. Nat. Commun. 7, 12881 (2016).

20. D. Shu, J. Guo, B. Zhang, Y. He, G. Wei, rDNA- and rRNA-derived communities present divergent assemblage patterns and functional traits throughout full-scale landfill leachate treatment process trains. Sci. Total Environ. 646, 1069–1079 (2019).

21. N. I. Wisnoski, M. E. Muscarella, M. L. Larsen, A. L. Peralta, J. T. Lennon, Metabolic insight into bacterial community assembly across ecosystem boundaries. Ecology. 101, e02968 (2020).

22. J. C. Stegen, X. Lin, A. E. Konopka, J. K. Fredrickson, Stochastic and deterministic assembly processes in subsurface microbial communities. ISME J. 6, 1653–1664 (2012).

23. E. B. Graham, J. C. Stegen, Dispersal-Based Microbial Community Assembly Decreases Biogeochemical Function. Processes. 5, 65 (2017).

24. F. Dini-Andreote, J. C. Stegen, J. D. van Elsas, J. F. Salles, Disentangling mechanisms that mediate the balance between stochastic and deterministic processes in microbial succession. Proc. Natl. Acad. Sci. 112, E1326–E1332 (2015).

25. J. C. Stegen, X. Lin, J. K. Fredrickson, A. E. Konopka, Estimating and mapping ecological processes influencing microbial community assembly. Front. Microbiol. 6 (2015), doi:10.3389/fmicb.2015.00370.

26. J. C. Stegen, X. Lin, J. K. Fredrickson, X. Chen, D. W. Kennedy, C. J. Murray, M. L. Rockhold, A. Konopka, Quantifying community assembly processes and identifying features that impose them. ISME J. 7, 2069–2079 (2013).

27. J. Zhou, D. Ning, Stochastic Community Assembly: Does It Matter in Microbial Ecology? Microbiol. Mol. Biol. Rev. 81 (2017), doi:10.1128/MMBR.00002-17.

28. J. Grilli, G. Barabás, M. J. Michalska-Smith, S. Allesina, Higher-order interactions stabilize dynamics in competitive network models. Nature. 548, 210–213 (2017).

29. E. M. Bottos, D. W. Kennedy, E. B. Romero, S. J. Fansler, J. M. Brown, L. M. Bramer, R. K. Chu, M. M. Tfaily, J. K. Jansson, J. C. Stegen, Dispersal limitation and thermodynamic constraints govern spatial structure of permafrost microbial communities. FEMS Microbiol. Ecol. 94 (2018), doi:10.1093/femsec/fiy110.

30. Y. Feng, R. Chen, J. C. Stegen, Z. Guo, J. Zhang, Z. Li, X. Lin, Two key features influencing community assembly processes at regional scale: Initial state and degree of change in environmental conditions. Mol. Ecol. 27, 5238–5251 (2018).

31. S. D. Jurburg, I. Nunes, J. C. Stegen, X. Le Roux, A. Priemé, S. J. Sørensen, J. F. Salles, Autogenic succession and deterministic recovery following disturbance in soil bacterial communities. Sci. Rep. 7, 1–11 (2017).

32. A. Sengupta, J. Indivero, C. Gunn, M. M. Tfaily, R. K. Chu, J. Toyoda, V. L. Bailey, N. D. Ward, J. C. Stegen, Spatial gradients in the characteristics of soil-carbon fractions are associated with abiotic features but not microbial communities. Biogeosciences. 16, 3911–3928 (2019).

33. E. B. Graham, M. M. Tfaily, A. R. Crump, A. E. Goldman, L. M. Bramer, E. Arntzen, E. Romero, C. T. Resch, D. W. Kennedy, J. C. Stegen, Carbon Inputs From Riparian Vegetation Limit Oxidation of Physically Bound Organic Carbon Via Biochemical and Thermodynamic Processes. J. Geophys. Res. Biogeosciences. 122, 3188–3205 (2017).

34. J. C. Stegen, T. Johnson, J. K. Fredrickson, M. J. Wilkins, A. E. Konopka, W. C. Nelson, E. V. Arntzen, W. B. Chrisler, R. K. Chu, S. J. Fansler, E. B. Graham, D. W. Kennedy, C. T. Resch, M. Tfaily, J. Zachara, Influences of organic carbon speciation on hyporheic corridor biogeochemistry and microbial ecology. Nat. Commun. 9, 1–11 (2018).

35. J. C. Stegen, A. Konopka, J. P. McKinley, C. Murray, X. Lin, M. D. Miller, D. W. Kennedy, E. A. Miller, C. T. Resch, J. K. Fredrickson, Coupling among Microbial Communities, Biogeochemistry and Mineralogy across Biogeochemical Facies. Sci. Rep. 6, 1–14 (2016).

36. P. Starnawski, T. Bataillon, T. J. G. Ettema, L. M. Jochum, L. Schreiber, X. Chen, M. A. Lever, M. F. Polz, B. B. Jørgensen, A. Schramm, K. U. Kjeldsen, Microbial community assembly and evolution in subseafloor sediment. Proc. Natl. Acad. Sci. 114, 2940–2945 (2017).

37. W. Wu, H.-P. Lu, A. Sastri, Y.-C. Yeh, G.-C. Gong, W.-C. Chou, C.-H. Hsieh, Contrasting the relative importance of species sorting and dispersal limitation in shaping marine bacterial versus protist communities. ISME J. 12, 485–494 (2018).

38. W. Chen, K. Ren, A. Isabwe, H. Chen, M. Liu, J. Yang, Stochastic processes shape microeukaryotic community assembly in a subtropical river across wet and dry seasons. Microbiome. 7, 138 (2019).

39. I. Martínez, J. C. Stegen, M. X. Maldonado-Gómez, A. M. Eren, P. M. Siba, A. R. Greenhill, J. Walter, The Gut Microbiota of Rural Papua New Guineans: Composition, Diversity Patterns, and Ecological Processes. Cell Rep. 11, 527–538 (2015).

40. I. D. Ofiţeru, M. Lunn, T. P. Curtis, G. F. Wells, C. S. Criddle, C. A. Francis, W. T. Sloan, Combined niche and neutral effects in a microbial wastewater treatment community. Proc. Natl. Acad. Sci. 107, 15345–15350 (2010).

41. J. Zhou, W. Liu, Y. Deng, Y.-H. Jiang, K. Xue, Z. He, J. D. V. Nostrand, L. Wu, Y. Yang, A. Wang, Stochastic Assembly Leads to Alternative Communities with Distinct Functions in a Bioreactor Microbial Community. mBio. 4 (2013), doi:10.1128/mBio.00584-12.

42. X. Jia, F. Dini-Andreote, J. Falcão Salles, Comparing the Influence of Assembly Processes Governing Bacterial Community Succession Based on DNA and RNA Data. Microorganisms. 8, 798 (2020).

43. R. A. Daly, M. A. Borton, M. J. Wilkins, D. W. Hoyt, D. J. Kountz, R. A. Wolfe, S. A. Welch, D. N. Marcus, R. V. Trexler, J. D. MacRae, J. A. Krzycki, D. R. Cole, P. J. Mouser, K. C. Wrighton, Microbial metabolisms in a 2.5-km-deep ecosystem created by hydraulic fracturing in shales. Nat. Microbiol. 1, 1–9 (2016).

44. G. E. Leventhal, M. Ackermann, K. T. Schiessl, Why microbes secrete molecules to modify their environment: the case of iron-chelating siderophores. J. R. Soc. Interface. 16, 20180674 (2019).

45. C. Ratzke, J. Denk, J. Gore, Ecological suicide in microbes. Nat. Ecol. Evol. 2, 867–872 (2018).

46. J. C. Stegen, E. M. Bottos, J. K. Jansson, A unified conceptual framework for prediction and control of microbiomes. Curr. Opin. Microbiol. (2018), doi:10.1016/j.mib.2018.06.002.

47. E. B. Graham, A. R. Crump, D. W. Kennedy, E. Arntzen, S. Fansler, S. O. Purvine, C. D. Nicora, W. Nelson, M. M. Tfaily, J. C. Stegen, Multi’omics comparison reveals metabolome biochemistry, not microbiome composition or gene expression, corresponds to elevated biogeochemical function in the hyporheic zone. Sci. Total Environ. 642, 742–753 (2018).

48. K. Boye, V. Noël, M. M. Tfaily, S. E. Bone, K. H. Williams, J. R. Bargar, S. Fendorf, Thermodynamically controlled preservation of organic carbon in floodplains. Nat. Geosci. 10, 415–419 (2017).

49. V. A. Garayburu-Caruso, J. C. Stegen, H.-S. Song, L. Renteria, J. Wells, W. Garcia, C. T. Resch, A. E. Goldman, R. K. Chu, J. Toyoda, E. B. Graham, Carbon Limitation Leads to Thermodynamic Regulation of Aerobic Metabolism. Environ. Sci. Technol. Lett. (2020), doi:10.1021/acs.estlett.0c00258.

50. H.-S. Song, J. C. Stegen, E. B. Graham, J.-Y. Lee, V. A. Garayburu-Caruso, W. C. Nelson, X. Chen, J. D. Moulton, T. D. Scheibe, bioRxiv, in press, doi:10.1101/2020.02.27.968669.

51. M. E. McClain, E. W. Boyer, C. L. Dent, S. E. Gergel, N. B. Grimm, P. M. Groffman, S. C. Hart, J. W. Harvey, C. A. Johnston, E. Mayorga, W. H. McDowell, G. Pinay, Biogeochemical Hot Spots and Hot Moments at the Interface of Terrestrial and Aquatic Ecosystems. Ecosystems. 6, 301–312 (2003).

52. E. S. Bernhardt, J. R. Blaszczak, C. D. Ficken, M. L. Fork, K. E. Kaiser, E. C. Seybold, Control Points in Ecosystems: Moving Beyond the Hot Spot Hot Moment Concept. Ecosystems. 20, 665–682 (2017).

53. B. Arora, M. A. Briggs, J. Zarnetske, J. C. Stegen, J. Gomez-Velez, D. Dwivedi, C. I. Steefel, in Biogeochemistry of the Critical Zone, A. Wymore, W. Yang, W. Silver, B. McDowell, J. Chorover, Eds. (Springer-Nature, 2020).

54. F. Boano, J. W. Harvey, A. Marion, A. I. Packman, R. Revelli, L. Ridolfi, A. Wörman, Hyporheic flow and transport processes: Mechanisms, models, and biogeochemical implications. Rev. Geophys. 52, 603–679 (2014).

55. R. M. Burrows, H. Rutlidge, N. R. Bond, S. M. Eberhard, A. Auhl, M. S. Andersen, D. G. Valdez, M. J. Kennard, High rates of organic carbon processing in the hyporheic zone of intermittent streams. Sci. Rep. 7, 1–11 (2017).

56. B. O. L. Demars, Hydrological pulses and burning of dissolved organic carbon by stream respiration. Limnol. Oceanogr. 64, 406–421 (2019).

57. H. Fischer, F. Kloep, S. Wilzcek, M. T. Pusch, A River’s Liver – Microbial Processes within the Hyporheic Zone of a Large Lowland River. Biogeochemistry. 76, 349–371 (2005).

58. M. H. Kaufman, M. B. Cardenas, J. Buttles, A. J. Kessler, P. L. M. Cook, Hyporheic hot moments: Dissolved oxygen dynamics in the hyporheic zone in response to surface flow perturbations. Water Resour. Res. 53, 6642–6662 (2017).

59. S. T. Larned, T. Datry, D. B. Arscott, K. Tockner, Emerging concepts in temporary-river ecology. Freshw. Biol. 55, 717–738 (2010).

60. A. M. Romaní, E. Vázquez, A. Butturini, Microbial Availability and Size Fractionation of Dissolved Organic Carbon After Drought in an Intermittent Stream: Biogeochemical Link Across the Stream–Riparian Interface. Microb. Ecol. 52, 501–512 (2006).

61. H. F. Birch, Mineralisation of plant nitrogen following alternate wet and dry conditions. Plant Soil. 20, 43–49 (1964).

62. A. E. Goldman, E. B. Graham, A. R. Crump, D. W. Kennedy, E. B. Romero, C. G. Anderson, K. L. Dana, C. T. Resch, J. K. Fredrickson, J. C. Stegen, Biogeochemical cycling at the aquatic–terrestrial interface is linked to parafluvial hyporheic zone inundation history. Biogeosciences. 14, 4229–4241 (2017).

63. E. B. Graham, A. R. Crump, C. T. Resch, S. Fansler, E. Arntzen, D. W. Kennedy, J. K. Fredrickson, J. C. Stegen, Coupling Spatiotemporal Community Assembly Processes to Changes in Microbial Metabolism. Front. Microbiol. 7 (2016), doi:10.3389/fmicb.2016.01949.

64. J. M. Zachara, P. E. Long, J. Bargar, J. A. Davis, P. Fox, J. K. Fredrickson, M. D. Freshley, E. Konopka, C. Liu, J. P. McKinley, M. L. Rockhold, K. H. Williams, S. B. Yabusaki, Persistence of uranium groundwater plumes: Contrasting mechanisms at two DOE sites in the groundwater–river interaction zone. J. Contam. Hydrol. 147, 45–72 (2013).

65. L. D. Slater, D. Ntarlagiannis, F. D. Day-Lewis, K. Mwakanyamale, R. J. Versteeg, A. Ward, C. Strickland, C. D. Johnson, J. W. Lane, Use of electrical imaging and distributed temperature sensing methods to characterize surface water–groundwater exchange regulating uranium transport at the Hanford 300 Area, Washington. Water Resour. Res. 46 (2010), doi:10.1029/2010WR009110.

66. E. Arntzen, Effects of fluctuating river flow on groundwater/surface water mixing in the hyporheic zone of a regulated, large cobble bed river - Arntzen - 2006 - River Research and Applications - Wiley Online Library (2006), (available at https://onlinelibrary.wiley.com/doi/abs/10.1002/rra.947).

67. S. Fatichi, S. Manzoni, D. Or, A. Paschalis, A Mechanistic Model of Microbially Mediated Soil Biogeochemical Processes: A Reality Check. Glob. Biogeochem. Cycles. 33, 620–648 (2019).

68. E. B. Graham, A. R. Crump, C. T. Resch, S. Fansler, E. Arntzen, D. W. Kennedy, J. K. Fredrickson, J. C. Stegen, Deterministic influences exceed dispersal effects on hydrologically-connected microbiomes. Environ. Microbiol. 19, 1552–1567 (2017).

69. J. Brown, N. Zavoshy, C. J. Brislawn, L. A. McCue, “Hundo: a Snakemake workflow for microbial community sequence data” (e27272v1, PeerJ Inc., 2018), , doi:10.7287/peerj.preprints.27272v1.

70. R. E. Danczak, A. E. Goldman, R. K. Chu, J. G. Toyoda, V. A. Garayburu-Caruso, N. Tolić, E. B. Graham, J. W. Morad, L. Renteria, J. R. Wells, S. P. Herzog, A. S. Ward, J. C. Stegen, bioRxiv, in press, doi:10.1101/2020.02.12.946459.

71. D. E. LaRowe, P. Van Cappellen, Degradation of natural organic matter: A thermodynamic analysis. Geochim. Cosmochim. Acta. 75, 2030–2042 (2011).

72. B. M. Tripathi, J. C. Stegen, M. Kim, K. Dong, J. M. Adams, Y. K. Lee, Soil pH mediates the balance between stochastic and deterministic assembly of bacteria. ISME J. 12, 1072–1083 (2018).

73. J. Wang, J. Shen, Y. Wu, C. Tu, J. Soininen, J. C. Stegen, J. He, X. Liu, L. Zhang, E. Zhang, Phylogenetic beta diversity in bacterial assemblages across ecosystems: deterministic versus stochastic processes. ISME J. 7, 1310–1321 (2013).

74. L. Fillinger, Y. Zhou, C. Kellermann, C. Griebler, Non-random processes determine the colonization of groundwater sediments by microbial communities in a pristine porous aquifer. Environ. Microbiol. 21, 327–342 (2019).

75. Y. Li, Y. Gao, W. Zhang, C. Wang, P. Wang, L. Niu, H. Wu, Homogeneous selection dominates the microbial community assembly in the sediment of the Three Gorges Reservoir. Sci. Total Environ. 690, 50–60 (2019).

76. A. Sengupta, J. C. Stegen, A. A. M. Neto, Y. Wang, J. W. Neilson, T. Tatarin, E. Hunt, K. Dontsova, J. Chorover, P. A. Troch, R. M. Maier, Assessing Microbial Community Patterns During Incipient Soil Formation From Basalt. J. Geophys. Res. Biogeosciences. 124, 941–958 (2019).

77. T. Whitman, R. Neurath, A. Perera, I. Chu-Jacoby, D. Ning, J. Zhou, P. Nico, J. Pett-Ridge, M. Firestone, Microbial community assembly differs across minerals in a rhizosphere microcosm. Environ. Microbiol. 20, 4444–4460 (2018).

78. E. S. Boyd, D. E. Cummings, G. G. Geesey, Mineralogy Influences Structure and Diversity of Bacterial Communities Associated with Geological Substrata in a Pristine Aquifer. Microb. Ecol. 54, 170–182 (2007).

79. J. K. Carson, L. Campbell, D. Rooney, N. Clipson, D. B. Gleeson, Minerals in soil select distinct bacterial communities in their microhabitats. FEMS Microbiol. Ecol. 67, 381–388 (2009).

80. S. Doetterl, A. A. Berhe, C. Arnold, S. Bodé, P. Fiener, P. Finke, L. Fuchslueger, M. Griepentrog, J. W. Harden, E. Nadeu, J. Schnecker, J. Six, S. Trumbore, K. Van Oost, C. Vogel, P. Boeckx, Links among warming, carbon and microbial dynamics mediated by soil mineral weathering. Nat. Geosci. 11, 589–593 (2018).

81. B. Fauvel, H.-M. Cauchie, C. Gantzer, L. Ogorzaly, Influence of physico-chemical characteristics of sediment on the in situ spatial distribution of F-specific RNA phages in the riverbed. FEMS Microbiol. Ecol. 95 (2019), doi:10.1093/femsec/fiy240.

82. B. S. Mauck, J. A. Roberts, Mineralogic Control on Abundance and Diversity of Surface-Adherent Microbial Communities. Geomicrobiol. J. 24, 167–177 (2007).

83. Z. B. Freedman, K. J. Romanowicz, R. A. Upchurch, D. R. Zak, Differential responses of total and active soil microbial communities to long-term experimental N deposition. Soil Biol. Biochem. 90, 275–282 (2015).

84. D. J. Levy-Booth, I. J. W. Giesbrecht, C. T. E. Kellogg, T. J. Heger, D. V. D’Amore, P. J. Keeling, S. J. Hallam, W. W. Mohn, Seasonal and ecohydrological regulation of active microbial populations involved in DOC, CO 2, and CH 4 fluxes in temperate rainforest soil. ISME J. 13, 950–963 (2019).

85. D. S. Baldwin, A. M. Mitchell, The effects of drying and re-flooding on the sediment and soilnutrient dynamics of lowland river–floodplain systems: a synthesis. Regul. Rivers Res. Manag. 16, 457–467 (2000).

86. S. Manzoni, J. P. Schimel, A. Porporato, Responses of soil microbial communities to water stress: results from a meta-analysis. Ecology. 93, 930–938 (2012).

87. S. Pérez Castro, E. E. Cleland, R. Wagner, R. A. Sawad, D. A. Lipson, Soil microbial responses to drought and exotic plants shift carbon metabolism. ISME J. 13, 1776–1787 (2019).

88. J. I. Prosser, J. B. H. Martiny, Conceptual challenges in microbial community ecology. Philos. Trans. R. Soc. B Biol. Sci. 375, 20190241 (2020).

89. H. F. Birch, M. T. Friend, Humus Decomposition in East African Soils. Nature. 178, 500–501 (1956).

90. N. Fierer, A. S. Allen, J. P. Schimel, P. A. Holden, Controls on microbial CO2 production: a comparison of surface and subsurface soil horizons. Glob. Change Biol. 9, 1322–1332 (2003).

91. G. Gionchetta, F. Oliva, A. M. Romani, L. Baneras, Hydrological variations shape diversity and functional responses of streambed microbes. Sci. Total Environ. 714, 136838 (2020).

92. P. M. Homyak, J. C. Blankinship, E. W. Slessarev, S. M. Schaeffer, S. Manzoni, J. P. Schimel, Effects of altered dry season length and plant inputs on soluble soil carbon. Ecology. 99, 2348–2362 (2018).

93. S. Louca, M. F. Polz, F. Mazel, M. B. N. Albright, J. A. Huber, M. I. O’Connor, M. Ackermann, A. S. Hahn, D. S. Srivastava, S. A. Crowe, M. Doebeli, L. W. Parfrey, Function and functional redundancy in microbial systems. Nat. Ecol. Evol. 2, 936–943 (2018).

94. J. G. Caporaso, EMP 16S Illumina Amplicon Protocol (2018), doi:10.17504/protocols.io.nuudeww.

